# Transneuronal delivery of designer-cytokine enables functional recovery after complete spinal cord injury

**DOI:** 10.1101/831271

**Authors:** Marco Leibinger, Charlotte Zeitler, Philipp Gobrecht, Anastasia Andreadaki, Dietmar Fischer

**Author notes:** Correspondence and proofs: Dietmar Fischer, Ph.D., Department of Cell Physiology, Ruhr University of Bochum, Universitätsstraße 150, 44780 Bochum, Germanym. **Lead Contact:** Further information and requests for resources and reagents should be directed to and will be fulfilled by the Lead Contact Dietmar Fischer.

## Abstract

Spinal cord injury (SCI) often causes severe and permanent disabilities. The current study uses a transneuronal approach to stimulate spinal cord regeneration by AAV-hyper-IL-6 (hIL-6) application after injury. While preinjury PTEN knockout in cortical motoneurons fails to improve functional recovery after complete spinal cord crush, a single, postinjury injection of hIL-6 into the sensorimotor cortex markedly promotes axon regeneration in the corticospinal and, remarkably, raphespinal tracts enabling significant locomotion recovery of both hindlimbs. Moreover, transduced cortical motoneurons directly innervate serotonergic neurons in both sides of the raphe nuclei equally, enabling the synaptic release of hIL-6 and the transneuronal stimulation of raphe neurons in the brain stem. Functional recovery depends on the regeneration of serotonergic neurons as their degeneration induced by a toxin abolishes the hIL-6-mediated recovery. Thus, the transneuronal application of highly potent cytokines enables functional regeneration by stimulating neurons in the deep brain stem that are otherwise challenging to access, yet highly relevant for functional recovery after SCI.

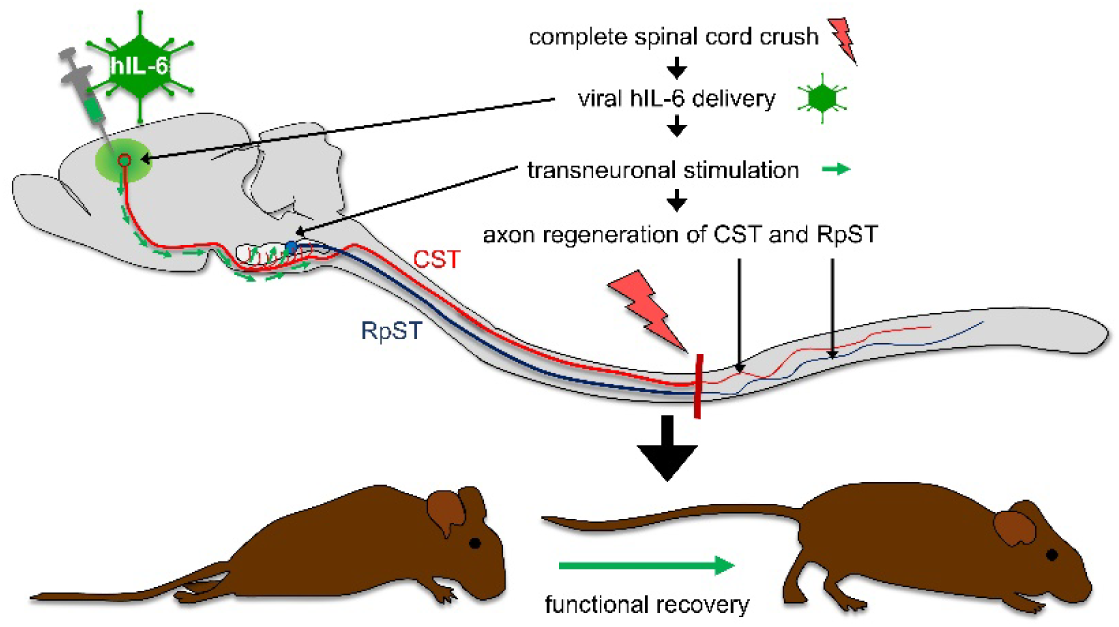

## Introduction

Neurons of the adult mammalian central nervous system (CNS) do not normally regenerate injured axons. This regenerative failure often causes severe and permanent disabilities, such as para- or tetraplegia after spinal cord injury. To date, no cures are available in the clinic, underscoring the need for novel therapeutic strategies enabling functional recovery in respective patients.

Besides an inhibitory environment for axonal growth cones caused by myelin or the forming glial scar, the lack of CNS regeneration is mainly attributed to a developmental decline in the neuron-intrinsic growth capacity of axons per se^1–3^. Among all descending pathways, the corticospinal tract (CST), which controls voluntary fine movements, is the most resistant to regeneration. Despite numerous efforts aiming to facilitate axon regrowth of the CST over the last decades, such as the delivery of neurotrophic factors^4–6^ or neutralizing inhibitory cues^7–9^, success has remained very limited. However, the conditional genetic knockout of the phosphatase and tensin homolog (PTEN^-/-^) in cortical motor neurons, which leads to an activation of the phosphatidylinositol-3-kinase/protein kinase B (PI3K/AKT)/mTOR signaling pathway, enables some regeneration of CST axons beyond the site of injury^10^. Although this approach facilitated the most robust anatomical regeneration of the CST after a complete spinal cord crush injury so far^10^, it fails to improve functional motor recovery^11^.

In the optic nerve, the activation of the Janus kinase/signal transducer and activator of transcription 3 (JAK/STAT3) pathway stimulates the regeneration of CNS axons^12, 13^. JAK/STAT3 activation is achieved via the delivery of IL-6-type cytokines such as CNTF, LIF, IL-6, and/or the genetic depletion of the intrinsic STAT3 feedback inhibitor: suppressor of cytokine signaling 3 (SOCS3)^12, 14–17^. However, the low and restricted expression of the cytokine-specific α-receptor subunits in CNS neurons that are required for signaling induction generally limits these pro-regenerative effects of native cytokines. For this reason, a gene therapeutic approach was recently developed using the designer cytokine hyper-interleukine-6 (hIL-6), which consists of the bioactive part of the IL-6 protein covalently linked to the soluble IL-6 receptor α subunit. In contrast to native cytokines, hIL-6 can directly bind to the signal-transducing receptor subunit glycoprotein 130 (GP130) abundantly expressed by almost all neurons and thereby circumvent the limitation of natural cytokines^19, 20^. Hyper-IL-6 is reportedly as potent as CNTF but activates cytokine-dependent signaling pathways significantly stronger in different types of neurons because of its higher efficacy^18^. In the visual system, virus-assisted gene therapy with hIL-6, even when applied only once post-injury, induces stronger optic nerve regeneration than a pre-injury induced PTEN knockout (PTEN^-/-^)^18^. Hence, this treatment is the most effective approach to stimulate optic nerve regeneration when applied after injury.

The current study analyzed the effect of cortically applied AAV-hIL-6 alone or in combination with PTEN^-/-^ on functional recovery after complete spinal cord crush (SCC). A single unilateral injection of AAV-hIL-6 applied after complete SCC into the sensorimotor cortex promoted regeneration of CST-axons stronger than PTEN^-/-^, and, remarkably, also of serotonergic fibers of the raphespinal tract, which enabled locomotion recovery of both hindlimbs. Moreover, this study shows that cortical motoneurons form synapses with raphe neurons deep in the brain stem allowing the axonal transport and synaptic release of hIL-6 to stimulate regeneration of serotonergic axons. Thus, transneuronal stimulation of neurons located deep in the brain stem using highly potent molecules might be a promising strategy to achieve functional repair in the injured or diseased human CNS.

## Results

### Hyper-IL-6 and PTEN knockout activate different signaling pathways

To investigate the impact of hIL-6 and PTEN^-/-^ on regeneration associated signaling pathways in motoneurons, we applied either AAV2-hIL-6 (coexpressing GFP), AAV2-Cre, or AAV2-GFP into the sensorimotor cortex of adult PTEN-floxed mice. As determined by GFP co-expression, the transduction was restricted to cortical cells, mainly motoneurons, adjacent to the injection sites (Fig. 1 A, B, E, G I; Fig. S1). In contrast to AAV2-Cre (PTEN^-/-^) and AAV2-GFP, AAV2-hIL-6 application induced strong STAT3-phosphorylation, as shown in sections of cortical tissue and western blot lysates (Fig. 1 A-C, K, L, J). STAT3 activation was already maximal after 1 week and remained stable over at least 8 weeks (Fig. 1 K, I). In contrast to PTEN^-/-^, which induced robust phosphorylation of AKT (pAKT) and S6 (pS6), AAV2-hIL-6 had little impact on the phosphorylation of these proteins (Fig. 1 K, M, N, P). Immunohistochemically stained cortical sections verified Western blot results (Fig. 1 A-J). Moreover, pSTAT3-positive signals in all GFP-positive hIL-6-transduced neurons and adjacent cells indicated the paracrine effects of released hIL-6 (Fig. 1 B, C). PTEN^-/-^-induced phosphorylation of S6 was restricted to GFP-positive (transduced) neurons only (Fig. 1 H, I). Neither hIL-6 nor PTEN^-/-^ influenced phosphorylation of ERK1/2 (Fig. 1 N, O).

**Figure 1:**
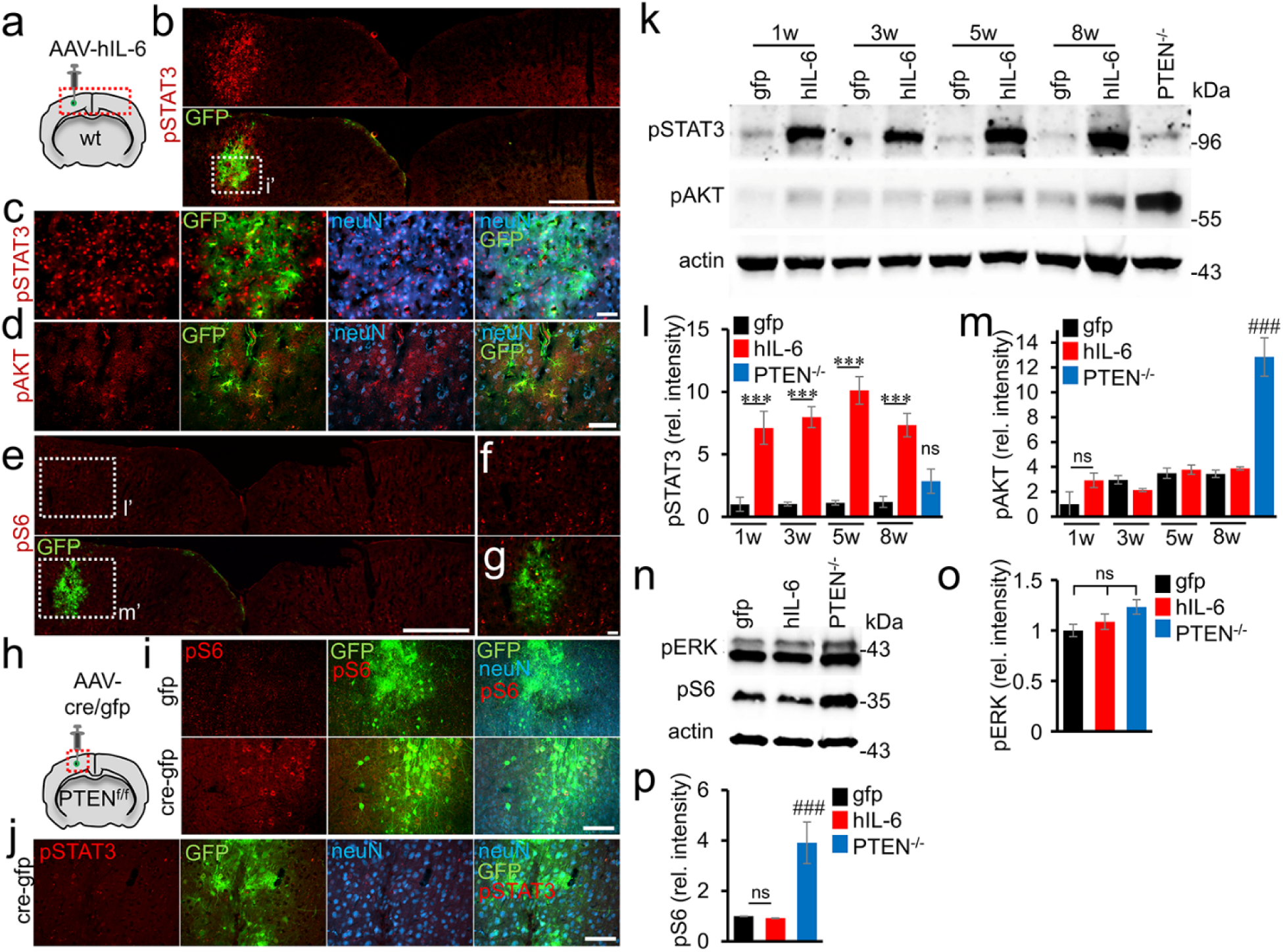
Signaling pathway activation by hIL-6 or PTEN^-/-^. **A)** Schematic drawing illustrating the location of the AAV-injection site (dashed box) shown in **H**. **B)** Coronal section of the sensorimotor cortex from a wild type mouse three weeks after intracortical injection of AAV2-hIL-6 into the left hemisphere. The section was immunohistochemically stained for phosphorylated STAT3 (pSTAT3, red). GFP (green) was co-expressed by the AAV2-hIL-6. Scale bar = 500 µm. **C)** Higher magnification of dotted box shown in **B** with additional blue channel showing neuron-specific nuclear staining (NeuN). Scale bar = 50 µm. **D-G)** Immunohistochemical staining against T308-phosphorylated AKT (pAKT, **D**, red) and phosphorylated ribosomal protein S6 (pS6, **E-G**, red) of sections described in A. Dashed boxes are shown at higher magnification in **F** and **G**. Scale bar = 50 µm (**D, F, G**); 500 µm (**E**). **H)** Schematic drawing illustrating the injection site and location of images shown in **I** and **J**. **I/J)** Cortical sections of PTEN^f/f^ mice <u>3</u> weeks after AAV2-Cre-GFP (Cre-<u>gfp</u>), or AAV2-GFP (<u>gfp</u>) injections, stained for pS6 (**I**, red), or pSTAT3 (**J**, red). Scale bar = 50 µm. **K)** Western blot analysis: Lysates prepared from the sensorimotor cortex of PTEN^f/f^ mice 1, 3, 5, or 8 weeks (w) after intracortical injection of either AAV2-hIL-6 (hIL-6), AAV2-GFP (gfp) or 5 weeks after AAV2-Cre application leading to PTEN^-/-^. AAV2-hIL-6 induced STAT3 phosphorylation (pSTAT3) at all tested time points while PTEN^-/-^ only caused AKT phosphorylation at T308. Beta-actin served as a loading control. **L/M)** Densitometric quantifications of western blots depicted in **K**. Values represent means +/- SEM of samples from 3-4 animals per group. **N)** Western blot analysis of cortical lysates: Phosphorylation of ERK1/2 (pERK) was not altered by AAV2-GFP-, AAV2-hIL-6, or AAV2-Cre (PTEN^-/-^) 5 weeks after intracortical application. Only PTEN^-/-^ induced significant S6-phosphorylation (pS6). **O/P)** Densitometric quantification of western blots depicted in **N**. Values represent means +/- SEM of 4 independent cortical lysates (n=4) per group. Significances of intergroup differences in **I, M, O, P** were evaluated using a one-way analysis of variance (ANOVA) followed by Tukey post hoc test. Treatment effects compared to <u>gfp</u> (week 8): ^###^p<0.001; or as indicated: ***p<0.001, ns=non-significant.

### Hyper-IL-6 promotes CST regeneration

Next, we tested whether AAV2-hIL-6 application, PTEN^-/-^, or their combination affect corticospinal tract (CST) regeneration following complete spinal cord crush (SCC). As AAV2 reaches higher neuronal transduction rates in newborn animals^10, 11^, and to keep the methodology comparable to these previous studies, PTEN floxed mice received injections of either AAV2-Cre (PTEN^-/-^) or AAV2-GFP (PTEN^+/+^) into the left sensorimotor cortex at postnatal day 1 (P1). After 7 weeks, mice were subjected to SCC, and each received a second intracortical injection of either AAV2-hIL-6 or AAV2-GFP immediately afterward (Fig. 2 A), resulting in four experimental groups: (a) Control animals that had received AAV2-GFP injections twice, (b) PTEN^-/-^-mice that were treated with AAV-Cre and later with AAV2-GFP, (c) hIL-6 mice that received AAV2-GFP and later, after SCC, AAV2-hIL-6 and (d) PTEN^-/-^/hIL-6 mice that were treated with AAV2-Cre first and, after SCC, with AAV2-hIL-6. Axonal biotinylated dextran amine (BDA)-tracing of the CST (Fig. S2 J) was performed 6 weeks after SCC, followed by the collection of the brains and spinal cords for imaging 2 weeks after that (Fig. 2 A).

**Figure 2:**
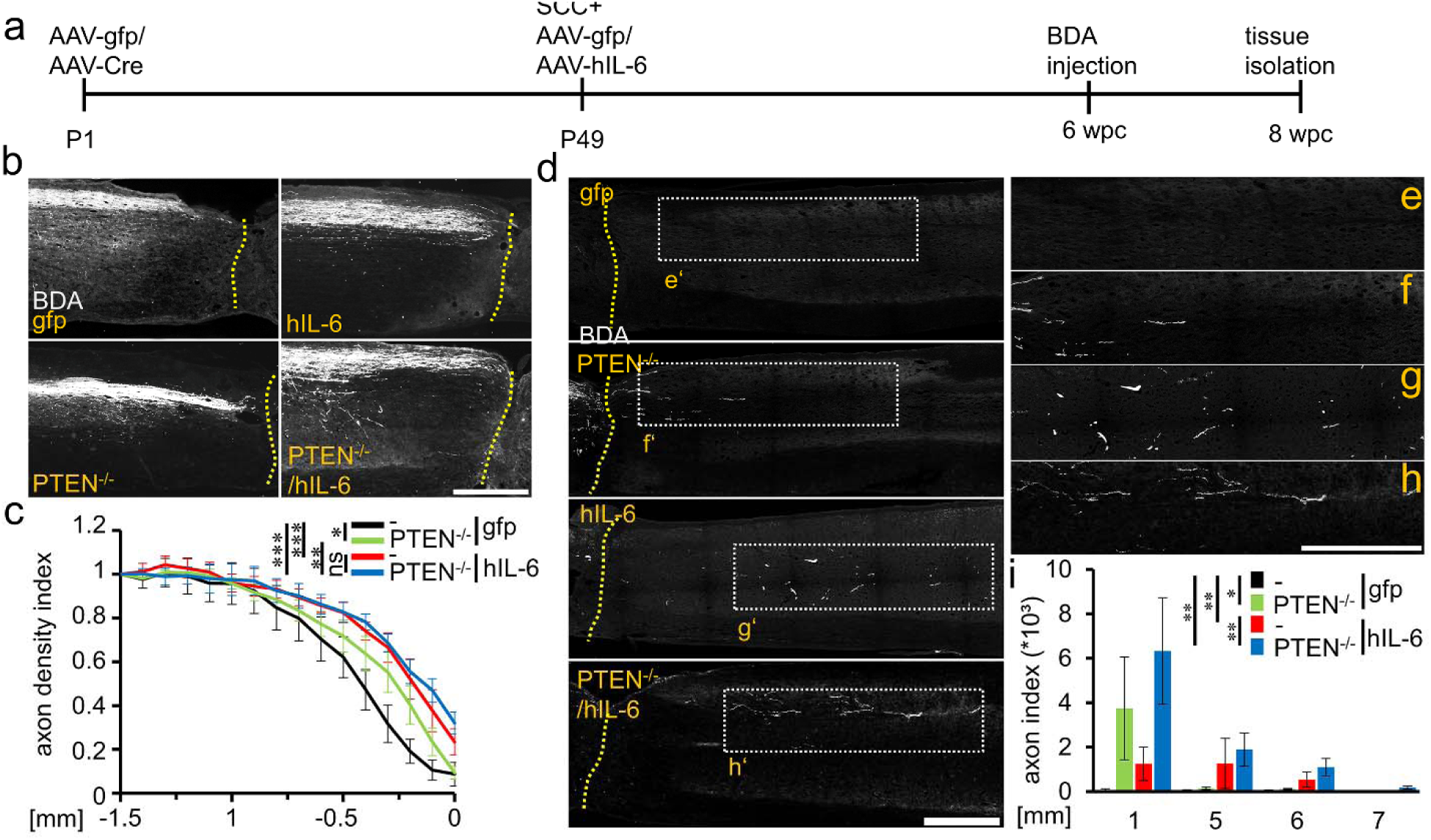
Hyper-IL-6 promotes CST axon regeneration after complete spinal cord crush. **A)** Timeline of surgical interventions for experiments presented in **B, C,** and Figs. 2-4. **B)** Sagittal thoracic spinal cord sections of four differently treated groups showing BDA-labeled axons (white) of the main CST rostral to the lesion site (dotted line) that retract or not. PTEN^f/f^ mice received injections of AAV2-GFP (PTEN^+/+^) or AAV2-Cre (PTEN^-/-^) at P1. After 7 weeks PTEN^+/+^ and PTEN^-/-^ mice were then subjected to spinal cord crush (SCC) (T8) and subsequently received intracortical injections of either AAV2-hIL-6 (hIL-6) or AAV2-GFP (gfp). Tissues were isolated 8 weeks after the SCC. Scale bar: 500 µm. **C)** Axon density index of CST fibers analyzing all spinal cord sections with the main CST of the animals described in **B.** Values at distances from 1.5 mm rostral to the lesion site (−1.5) up to the lesion core (0) were determined. Significances of intergroup differences were evaluated using a two-way analysis of variance (ANOVA) with Holm-Sidak post hoc test. Values represent means +/- SEM of 5-9 animals per group (PTEN^+/+^/gfp, n=5; PTEN^+/+^/hIL-6, n=9; PTEN^-/-^/gfp, n=6; PTEN^-/-^/hIL-6, n=9). Treatment effects as indicated: *p<0.05; **p<0.01; ***p<0.001; ns=non-significant. **D)** Representative images of BDA-labeled sagittal spinal cord sections from PTEN-floxed mice (PTEN^f/f^) after treatments as described in B. PTEN^f/f^ received intracortical injections of either AAV2-GFP (PTEN^+/+^) or AAV2-Cre (PTEN^-/-^) at P1 and were subjected to spinal cord crush (T8) 7 weeks later. Animals then received intracortical AAV2-GFP (gfp) or AAV2-hIL-6 (hIL-6) injections immediately afterward. Tissues were isolated after 8 weeks. BDA-labeled (white) regenerating axons of the central CST beyond the lesion site (dotted line) were seen only after PTEN^-/-^, hIL-6, or hIL-6/PTEN^-/-^-treatments. Scale bar: 500 µm. **E-H)** Higher magnification of regenerating axons in dotted boxes as indicated in **D**. Scale bar: 500 µm. **I)** Quantification of regenerating CST axons at the indicated distances caudal to the lesion site. Axon numbers were divided by the total number of BDA-labeled CST fibers in the medulla (axon index) from animals as described in **D**. Values represent the mean +/- SEM of 5-9 animals per group (PTEN^+/+^/gfp, n=5; PTEN^+/+^/hIL-6, n=9; PTEN^-/-^/gfp, n=6; PTEN^-/-^/hIL-6, n=9). Significances of intergroup differences were evaluated using a two-way analysis of variance (ANOVA) with Tukey post hoc tests. Treatment effects as indicated: ***p<0.001; ns=non-significant

Neither PTEN^-/-^, hIL-6 expression nor their combination affected the total number of BDA-positive CST axons in the medullary pyramid (Fig. S2 A, B, J) or CST-axonal sprouting in the thoracic spinal cord rostral to the lesion site compared to controls (Fig. S2 C-J). However, axonal dieback of CST-fibers above the injury site, typically seen in control mice, was significantly reduced by PTEN^-/-^ and slightly stronger by the AAV2-hIL-6 treatment (Fig. 2 B, C). The combination of PTEN^-/-^ + AAV2-hIL-6 showed no additional effect (Fig. 2 B, C).

We then analyzed fiber regeneration beyond the crush site. Contrary to controls (Fig. 2 D, E, I; Fig S3 A), PTEN^-/-^ enabled regeneration of some CST axons, most of which did, however, not exceed distances longer than 1.5 mm (Fig. 2 D, F, I; Fig. S3 A). Strikingly, AAV2-hIL-6 treatment resulted in stronger CST regeneration with the longest axon reaching up to 6 mm (Fig. 2 D, G, I; Fig. S3 A). This effect was slightly further increased by the combination of PTEN^-/-^ and AAV2-hIL-6 with the longest axons reaching >7 mm (Fig. 2 D, H, I; Fig. S3 A). No BDA labeled axons were detected in cross-sections >11 mm past the lesion site, thereby excluding spared axons and verifying the completeness of axonal injury in all animals.

### Hyper-IL-6 promotes functional recovery

Before tissue isolation and analysis, hindlimb movement had been analyzed in all four experimental groups using open-field locomotion tests according to the Basso Mouse Scale (BMS) over the postinjury period of 8 weeks^21^. Consistent with previous reports^21–23^, the BMS score dropped down to 0 in all animals 1 day after SCC, also indicating the completeness of the SCC (Fig. 3 A, B; Fig. S4 A-E). Over time, control animals developed only active ankle movements, including spasms (supplementary video 1) as described previously^21, 22^, resulting in an average final score of 2 (Fig. 3 A, B; Fig. S4 A; supplementary video 1). Despite the effect on CST-regeneration (Fig. 2 D, F, I), PTEN^-/-^ did not significantly improve the BMS score compared to AAV2-GFP treated controls (Fig. 3 A, B; Fig. S4 C; supplementary video 2). Strikingly, AAV2-hIL-6 treatment increased the score to 4-5 by restoring plantar stepping with full weight support through the hindlimb followed by lift-off, forward limb advancement, and reestablishment of weight support at initial contact in most of the animals (Fig. 3 A, B; Fig. S4 B; supplementary video 3). Combinatorial treatment (PTEN^-/-^ + AAV2-hIL-6) slightly enhanced this effect further (Fig. 3 A, B; Fig. S3 D, supplementary video 4), mainly seen in the BMS subscore (Fig. 3 E). One animal of this group even reached a coordinated movement of fore- and hindlimbs (score: 7) (Fig. 3 A, B; Fig. S4 D). Interestingly, despite the unilateral AAV2-hIL-6 injection into the left sensorimotor cortex only, both hindlimbs showed similar recovery (Fig. 3 A, C, D) and a bilateral application into the left and right side had no additional effect (Fig. 3 F; Fig. S4 E). Thus, in contrast to PTEN^-/-^, a single unilateral postinjury application of AAV2-hIL-6 into the sensorimotor cortex enabled locomotion of both hindlimbs after complete SCC.

**Figure 3:**
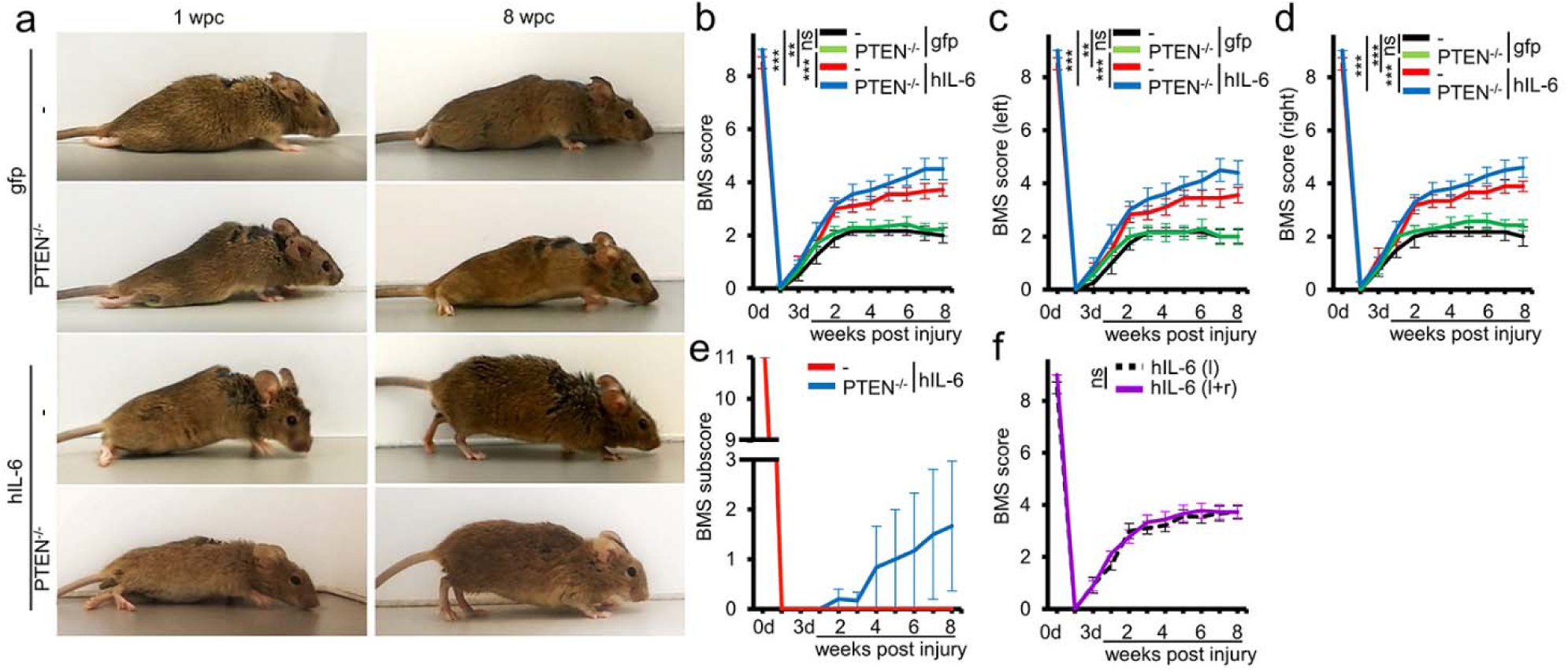
Hyper-IL-6 enables functional recovery after SCC. **A)** Representative pictures are showing open field hindlimb movement of mice at 1 and 8 weeks after spinal cord crush (wpc) and treatment, as described in Fig. 1. **B-D)** BMS score of animals as described in **A**, over eight weeks after spinal cord injury. Values represent means +/- SEM of 6-10 animals per group (PTEN^+/+^/gfp, n=6; PTEN^+/+^/hIL-6 n=9; PTEN^-/-^/gfp, n=7; PTEN^-/-^/hIL-6, n=10), showing either the average score of the left and right hind paw (**B**) or left (**C**) and right (**D**) side separately. **E)** BMS subscore of hIL-6 treated PTEN^+/+^(-) and PTEN^-/-^ mice as described in **A**. **F)** Average BMS score of left and right hind paws from mice after SCC and bilateral (left and right hemisphere (l+r)) intracortical injection of AAV2-hIL-6 compared to animals that had received a unilateral injection into the left hemisphere (l) only as presented in **B**. Values represent the mean +/- SEM of 9 animals per group (l, n=9; l+r, n=9). Significances of intergroup differences were evaluated using a two-way analysis of variance (ANOVA) with a Tukey post hoc test (**B-D**), or student’s t-test (**E, F**). Treatment effects as indicated: **p<0.01; ***p<0.001; ns=non-significant.

### Cortical AAV2-hIL-6 delivery promotes RpST regeneration

The lack of any functional recovery in PTEN^-/-^ mice suggested that improved CST regeneration was not the leading cause for the AAV2-hIL-6 mediated functional recovery. As descending serotonergic (5-HT-positive) axons of the raphespinal tract (RpST) are reportedly relevant for locomotor recovery^24–26^ we stained and analyzed these axons in sagittal spinal cord sections of the same animals used in the experiments before (Figs. 2-3). Control (AAV2-GFP) and PTEN^-/-^ mice revealed only sprouting of 5-HT-positive axons over distances of less than 1 mm beyond the crush site (Fig. 4 A-E, J; Fig. S3 C). Remarkably, AAV2-hIL-6 treated mice showed more and longer regeneration of serotonergic fibers than controls and PTEN^-/-^ (Fig. 4 A, F, G, J; Fig. S3 C) with the longest ones reaching >7 mm. Combinatorial treatment of AAV2-hIL-6 and PTEN^-/-^ had no additional effect (Fig. 4 A, H, I, J; Fig. S3 C). Furthermore, as measured for functional recovery (Fig. 3 F), bilateral AAV2-hIL-6 treatment did not increase RpST regeneration further compared to the unilateral virus application (Fig. 4 J; Fig. S3 C) and no 5-HT-positive axons were detected >11 mm beyond the injury site in any of these mice again verifying the absence of spared axons.

**Figure 4:**
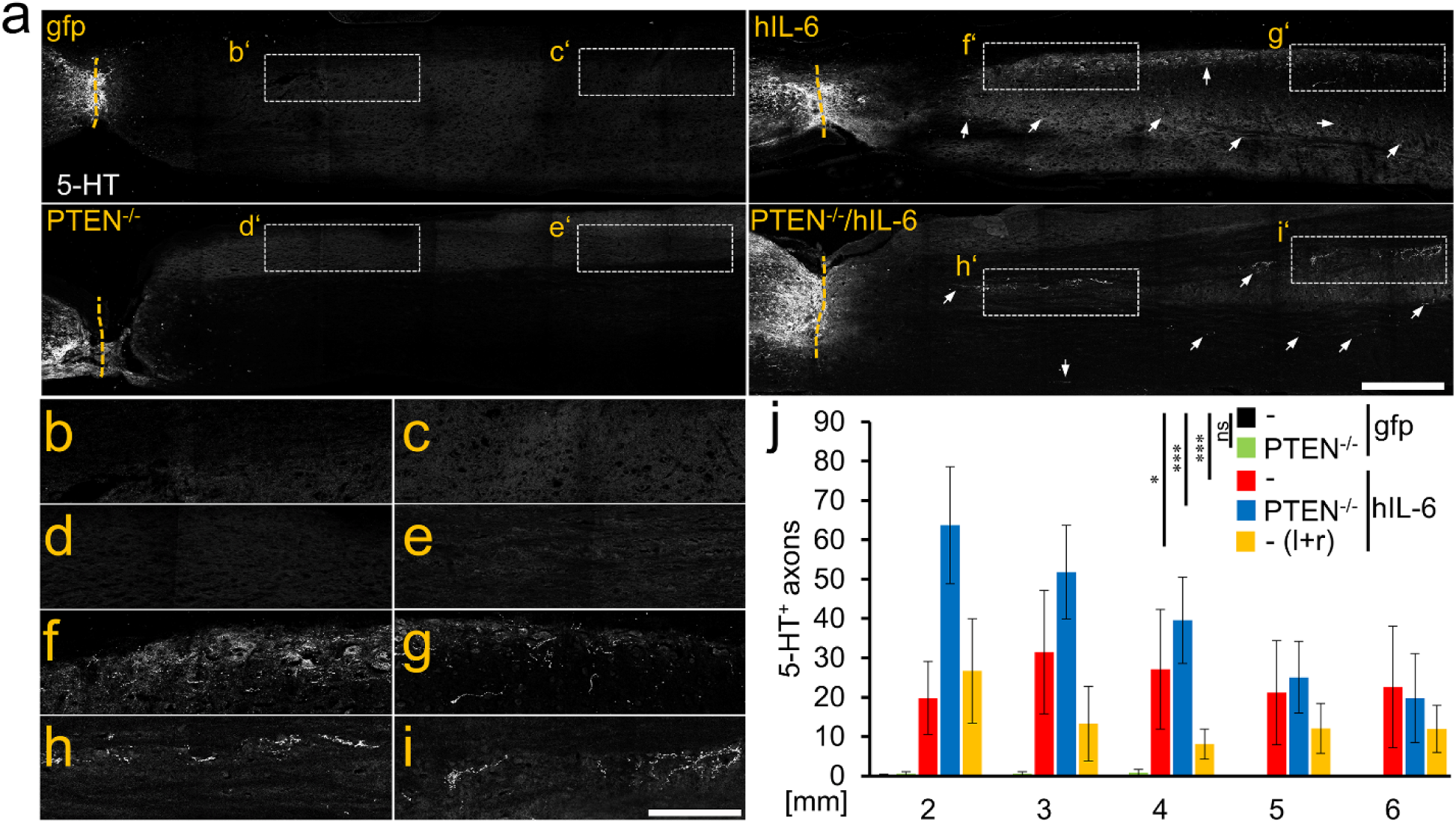
hIL-6 promotes axon regeneration of serotonergic fibers. **A)** Sagittal thoracic spinal cord sections isolated from PTEN^+/+^, or PTEN^-/-^ animals 8 weeks after spinal cord crush (SCC) and unilateral injection of AAV2-hIL-6 (hIL-6) or AAV2-GFP (gfp) (see Fig. 1). Raphe spinal tract (RpST) axons were stained using an anti-serotonin antibody (5-HT, white). Only AAV2-hIL-6-treated mice with or without additional PTEN^-/-^ showed significant regeneration of serotonergic axons beyond the lesion site (dashed line). Scale bar: 500 µm. **B-I)** Higher magnification of dashed boxes as indicated in **A**. Scale bar: 250 µm. **J)** Quantification of regenerating 5-HT-positive axons as described in **A** at indicated distances beyond the lesion. Values represent the mean +/- SEM of 5-10 animals per group (PTEN^+/+^/gfp, n=5; PTEN^+/+^/hIL-6, n=9; PTEN^-/-^/gfp, n=6; PTEN^-/-^/hIL-6, n=10; PTEN^+/+^/hIL-6 (l+r); n=5). Significances of intergroup differences were evaluated using a two-way analysis of variance (ANOVA) with a Holm Sidak post hoc test. Treatment effects as indicated: *p<0.05; **p<0.01; *** p<0.001; ns=non-significant.

### AAV2-hIL-6 mediated recovery depends on the regeneration of serotonergic neurons

To investigate the relevance of RpST regeneration for functional recovery, adult mice of the same genetic background were subjected to SCC and received bilateral intracortical injections of either AAV2-hIL-6 or AAV2-GFP directly afterward (Fig. 5 A). At week 6, when functional recovery in AAV2-hIL-6 treated mice had reached maximal levels, the neurotoxin 5, 7-Dihydroxytryptamine (DHT) was intracerebroventricularly injected to selectively kill serotonergic neurons (Fig. 5 A-D)^24, 27, 28^. One day after DHT application, the BMS in AAV2-hIL-6 treated mice dropped down to similar levels as in control animals, while DHT did not affect the BMS score of controls (Fig. 5 E, F; Supplementary video 5), suggesting a dependence of the AAV2-hIL-6 mediated locomotory recovery on RpST regeneration although AAV2-hIL-6 had not been applied into the brain stem where the serotonergic neurons originated.

**Figure 5:**
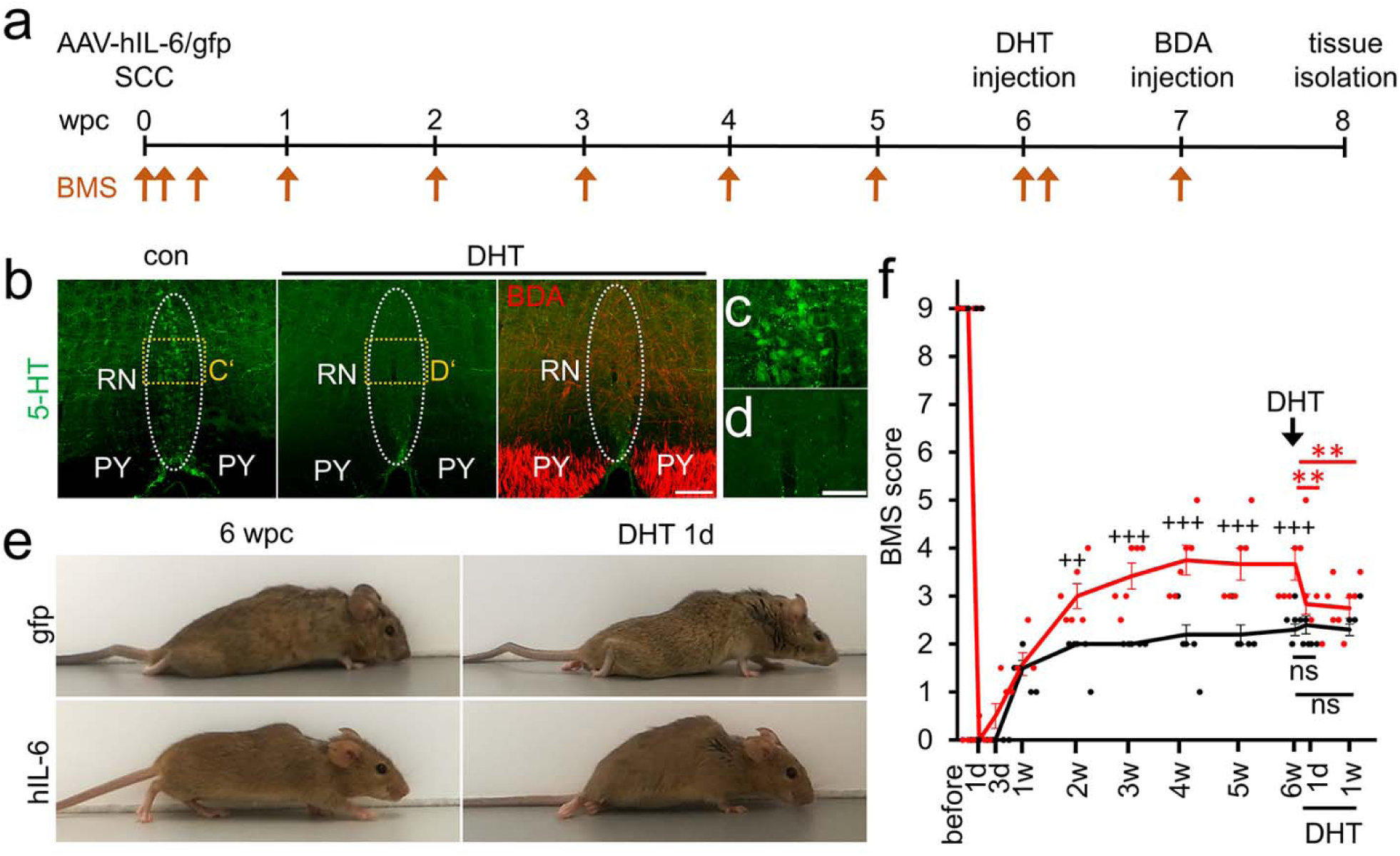
Regeneration of serotonergic axons is essential for functional recovery. **A)** Timeline of experiments shown in **B-F**. Adult mice were subjected to SCC and received bilateral intracortical AAV2-GFP (gfp) or AAV2-hIL-6 (hIL-6) injections. They were then monitored in open field movement and scored according to the Basso Mouse Scale (BMS) at the indicated time points (arrows) throughout 8 weeks post crush (wpc). Six weeks after SCC, the serotonin neurotoxin 5,7-dihydroxytrypatmine (DHT) was injected intracerebroventricularly into both hemispheres. One week before tissue isolation, BDA was injected intracortical to trace CST axons. **B)** Maximum intensity projection of confocal scans through 50 µm of cleared brain stem tissue from mice that received hIL-6 and DHT treatment as described in **A** compared to a control (con) without DHT treatment. Serotonergic neurons of raphe nuclei (RN, dotted line) were visualized by 5-HT staining (green). BDA-labeled CST axons in the pyramid (PY) are shown in red. DHT treatment eradicated all 5-HT-positive neurons without affecting corticospinal neurons. Scale bar: 100 µm **C/D)** Higher magnifications of dashed boxes (C’ and D’) in **B**. Scale bar: 50 µm **E)** Representative images of AAV2-GFP (gfp) or AAV2-hIL-6 (hIL-6)-treated mice 6 weeks post crush (wpc) before, and one day after DHT treatment. **F)** BMS score of animals as described in **A** at indicated time points after spinal cord crush and DHT treatment. Values represent means +/- SEM of 5-6 animals per group (gfp, n=5; hIL-6, n=6), showing the average score of left and right hind paws. Significances of intergroup differences were evaluated using a two-way analysis of variance (ANOVA) with a Holm Sidak post hoc test. Treatment effects of hIL-6 compared to gfp: ^++^p<0.01; ^+++^p<0.001; or as indicated: **p<0.01; ns=non-significant.

Since genetic backgrounds of mice could potentially affect regeneration, we also tested the postinjury applied AAV2-hIL-6 treatment in non-transgenic BL6 mice, whose low potential for functional recovery has been previously documented^21^. Eight weeks after complete SCC and unilateral intracortical AAV2-hIL-6 injection, BL-6 mice also showed reduced axonal dieback, but less CST-regeneration caudal to the lesion site than accordingly treated PTEN-floxed OLA mice (Fig. S5 A, B, Fig. S3 B). Nevertheless, the improvement in RpST axon regeneration and functional recovery were very similar in both mouse strains (Fig. S4 C-E; Fig. S3 D; Supplementary video 6), revealing a correlation between anatomical RpST regeneration and functional recovery.

### AAV2-hIL-6 confers transneuronal stimulation of serotonergic fiber regeneration

To explain the mechanism underlying the bilateral, RpST-dependent functional recovery by a single unilateral AAV2-hIL-6 application into the sensorimotor cortex, we tested the hypothesis that transduced motoneurons project axons to the raphe nuclei in the brain stem and that synaptically released hIL-6-protein stimulates regeneration of serotonergic neurons trans-synaptically. To this end, we initially used axon isolation devices to separate somata and axons of cultured sensory DRG neurons^29^ (Fig. S6 A) and transduced these neurons by adding baculoviruses (BV)^18, 30^ for either hIL-6 or GFP expression into the somal chamber only. Synaptotagmin-positive vesicles containing hIL-6 were found in the axons and their tips in the axonal chamber (Fig. S6 D). Additionally, Western blot analysis detected hIL-6 protein in the medium of the axon chamber of hIL-6 transduced neurons, verifying its release (Fig. S6 C). To test the functional activity of axonally released hIL-6, the medium of the axonal chamber was used to prepare dissociated cultures of adult retinal ganglion cells (RGC) (Fig. S6 B). Consistent with previous findings that hIL-6 promotes neurite extension of RGCs^18^, the conditioned medium from hIL-6 transduced sensory axons induced STAT3 phosphorylation (Fig. S6 E) and increased neurite outgrowth of RGCs twofold compared to the medium from GFP controls (Fig. S6 F, G). Thus, hIL-6 is transported in axons, released at the terminals, and remains active to stimulate axon regeneration.

To investigate whether transduced hIL-6 can also stimulate neurons trans-synaptically in vivo, we injected either AAV2-hIL-6 or AAV2-GFP into the left eye to transduce RGCs. Three weeks later, we tested for STAT3 phosphorylation in their brain targets, the lateral geniculate nucleus (LGN) and suprachiasmatic nucleus (SCN) (Fig. S6 G-I). In rodents, ∼95% of the optic nerve axons cross in the optic chiasm to the contralateral side. Consistent with a synaptic release of hIL-6, we found many pSTAT3-positive cellular nuclei near GFP-positive axon terminals of AAV2-hIL-6 transduced RGCs in the right hemisphere of LGN and SNC, much less in the respective left hemispheres and no signals at all in AAV2-GFP treated controls (Fig. S6 H, I, J). The hIL-6 expression in RGCs did not affect the integrity of axons as their number determined after neurofilament staining in the optic nerve and optic tract cross-sections was not reduced compared to untreated controls, excluding any uncontrolled or widespread release of the protein. We then investigated whether raphe nuclei in the brain stem receive synaptic input from transduced cortical motoneurons and analyzed 5-HT stained brain stem tissue from mice 8 weeks after SCC and AAV2-hIL-6 into the sensorimotor cortex as described above (Fig. 2 A). GFP-positive sprouts of cortical neurons were detected near 5-HT-positive raphe neurons (Fig. 6 A-D). Western blot analyses using brain stem lysates isolated 3 weeks after intracortical viral application showed pSTAT3 signals only in samples from AAV2-hIL-6-, but not AAV2-GFP-treated animals (Fig. 7 A, B). Additionally, transverse scans through cleared brain stem tissue from hIL-6 treated animals showed pSTAT3/5-HT double-positive neurons in the ipsi- and contralateral sides of the raphe nuclei in the medial column of the medulla (Fig. 7 C-D; Supplementary video 7). Similarly, coronal scans of cleared brain stem tissue of the nucleus raphe pallidus (NRPa) (Fig. 7 E, F) revealed pSTAT3-positive nuclei in serotonergic neurons after AAV2-hIL-6 but not AAV2-GFP treatment. On average, 46% (+/- 4.46 SEM) of the 5-HT-stained raphe neurons in AAV-2 hIL-6 treated animals were pSTAT3-positive. In contrast, AAV2-hIL-6 did not affect phosphorylation of either AKT or S6 in brain stem lysates compared to AAV2-GFP treated controls (Fig. 7 A, B). Moreover, intracortical AAV2-HIL-6 treatment also activated STAT3 in neurons of the red nucleus, a known target of cortical motoneurons (Fig. 7 G-I)^31^, thereby corroborating transneuronal delivery of hIL-6.

**Figure 6:**
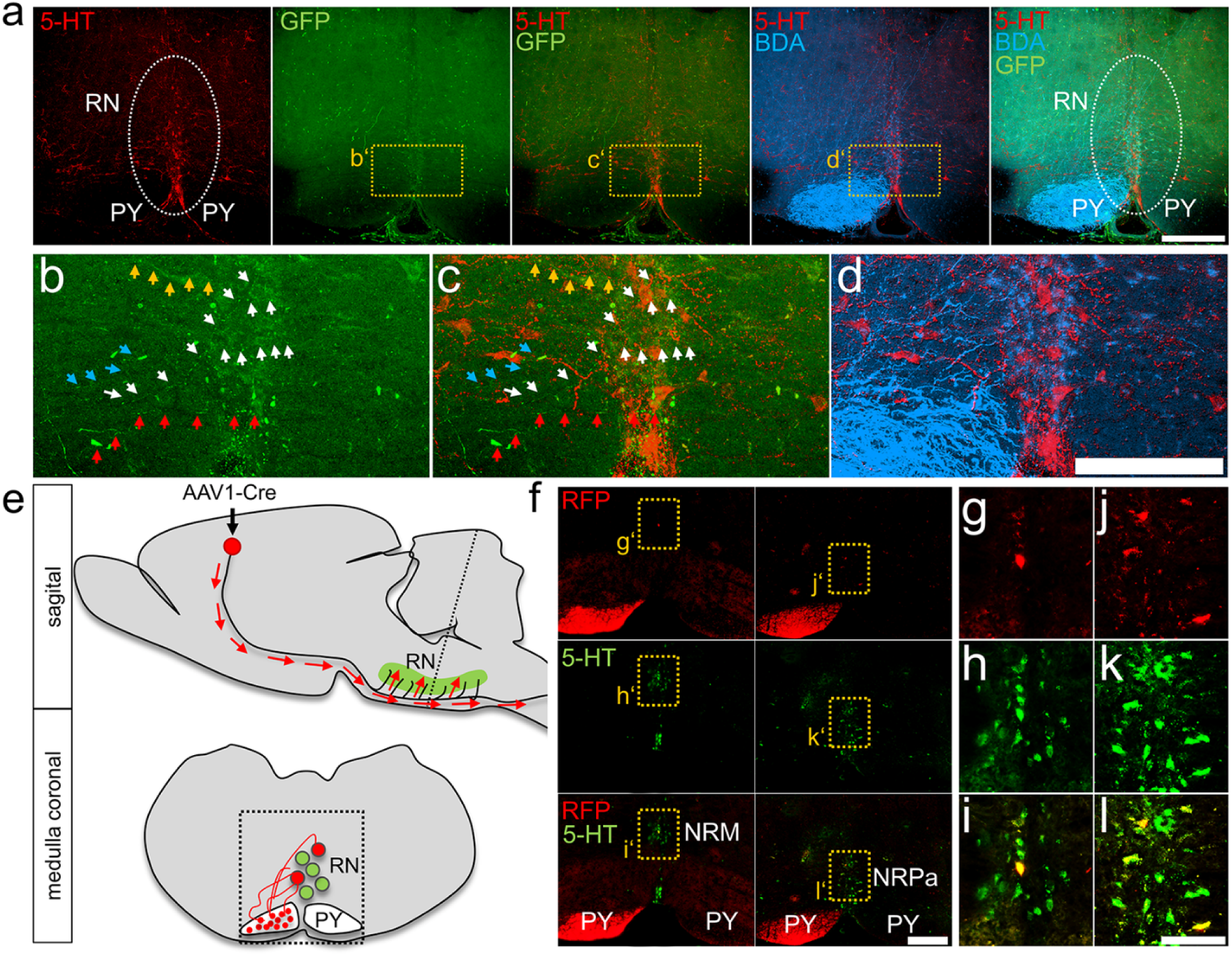
CST axon collaterals terminate in the brain stem raphe nuclei. **A)** Maximum intensity projection of coronal scan through 50 µm of cleared brain stem tissue from mice subjected to SCC and AAV2-hIL-6 injection as described in Fig. 1 **A:** BDA-traced pyramidal (PY) CST axons (blue) and serotonergic neurons of the raphe nuclei (RN) were visualized by 5-HT immunostaining (red). Axons of AAV2-hIL-6 transduced CST neurons were visualized by GFP co-expression (green). Scale bar: 100 µm **B-D)** Higher magnification of dashed boxes, as presented in **A**. GFP-positive axons are indicated by colored arrows. Scale bar: 100 µm. **E)** Schematic illustration of axonal transport and terminal release of AAV1-Cre into raphe nuclei (RN, green) of *Rosa-tdTomato* reporter mice in a sagittal view. The coronal view of the medulla illustrates RFP-positive pyramidal (PY) axon sprouts from AAV1-Cre transduced cortical neurons. Some pyramidal axons terminate at raphe neurons (green), leading to RFP expression in these cells after transneuronal transduction with AAV1-Cre. **F)** Coronal medullary sections from *Rosa-tdTomato* mice 2 weeks after intracortical AAV1-Cre injection as described in **E.** Transneuronally transduced serotonergic neurons of the nucleus raphe magnus (NRM), and nucleus raphe pallidus (NRPa) were identified by 5-HT staining (green) and RFP fluorescence (red). Scale bar: 250 µm **G-L**) Higher magnification of dashed yellow boxes as indicated in F. Scale bar: 100 µm

**Figure 7:**
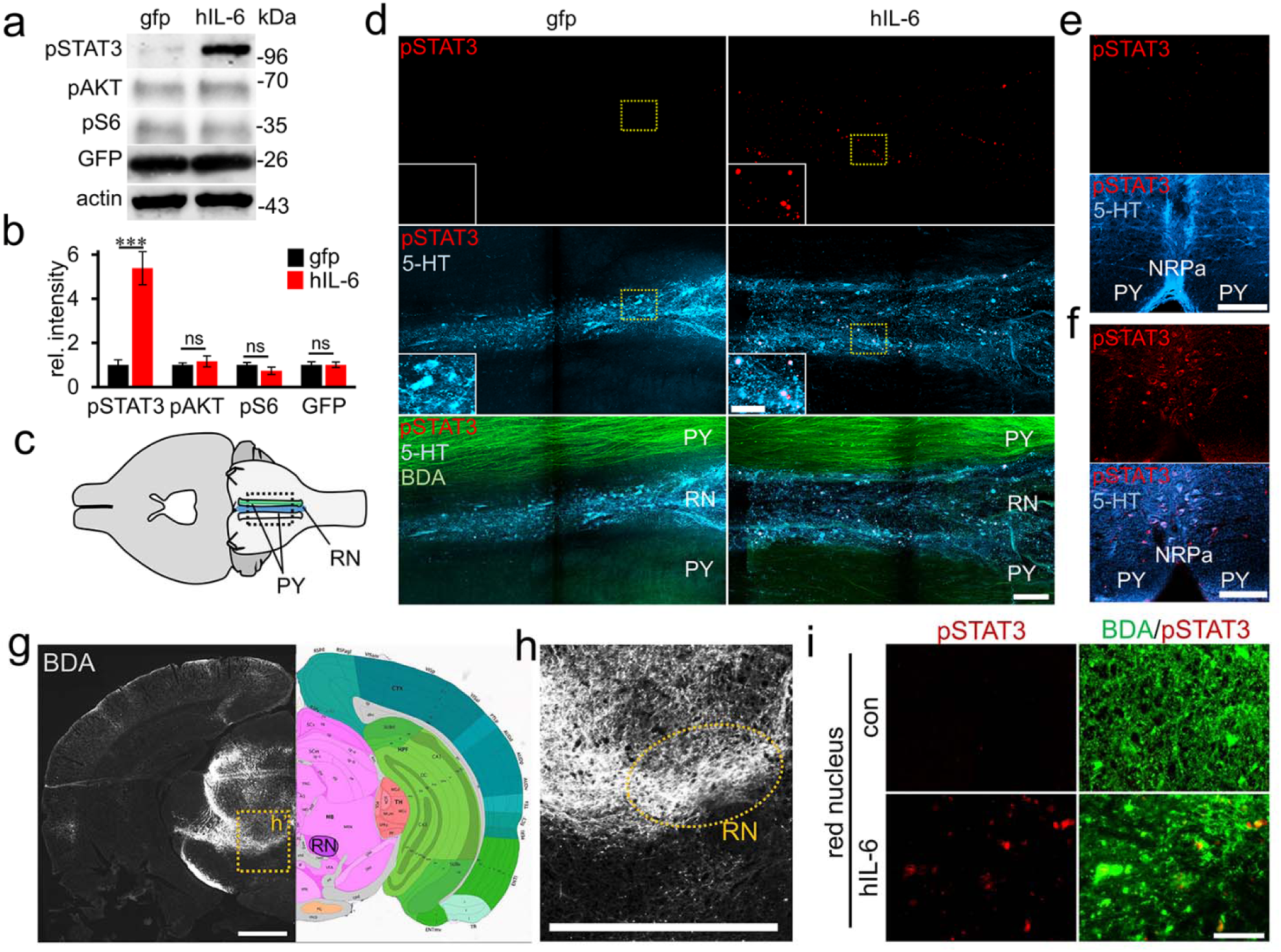
Hyper-IL-6 transneuronally stimulates neurons of raphe and red nuclei. **A)** Western blots: GFP and phosphorylated STAT3 (pSTAT3), AKT (pAKT), and S6 (pS6) were analyzed in lysates of the brain stem with raphe nuclei isolated 3 weeks after intracortical injection of either AAV2-GFP (gfp) or AAV2-hIL-6 (hIL-6). Similar GFP signals in lysates verified similar amounts of transduced collateral axons that projected to the brain stem. Beta-actin served as a loading control. **B)** Densitometric quantification of western blots from 3 different animals per group (n=3) after treatment as described in **A**. **C)** Schematic illustration related to images shown in **D:** Bottom view of the mouse brain showing pyramidal tracts (PY) and raphe nuclei (RN, blue) in the medial column of the medulla. Axons of GFP-expressing transduced cortical neurons in the left pyramid are indicated in green. The dotted box illustrates the area used for confocal scans from cleared tissue as shown in **D**. **D)** Maximum intensity projections of transverse confocal scans through 150 µm of cleared brain tissue as depictured in **C** from mice after unilateral (left) intracortical injections of AAV2-GFP (gfp) or AAV2-hIL-6 (hIL-6) and BDA treatment. Before tissue clearing, brains were stained against serotonin (5-HT, blue) and phosphorylated STAT3 (pSTAT3, red), while BDA labeled CST axons were visualized using fluorescently labeled streptavidin (BDA, green). Insets represent higher magnifications of areas in dashed yellow boxes. Scale bar: 50 µm, 25 µm for insets. **E/F)** Maximum intensity projections of coronal confocal scans through 50 µm of cleared brain stem tissue: PY and nucleus raphe pallidus (NRPa) of AAV2-GFP (**E**) and AAV2-hIL-6 (**F**) treated mice as described in **A.** Before clearing, tissues were stained for pSTAT3 (red) and serotonin (5-HT, blue) to identify raphe neurons. Scale bar: 100 µm Significances of intergroup differences in **B** were evaluated using the student’s t-test. Treatment effects compared to control: ***p<0.001; ns=non-significant. **G)** Coronal brain section 2 weeks after intracortical BDA injection showing BDA labeled axons of cortical motor neurons projecting into the red nucleus (RN). Localization of the RN is indicated by a corresponding map from Allen Brain Atlas (right). Scale bar 500 µm. **H)** Higher magnification of the dashed box shown in **G**. Scale bar: 500 µm **I)** Images of pSTAT3 (red) immunostained coronal sections of the ipsilateral red nucleus form mice 3 weeks after intracortical injection with either AAV2-GFP (con) or AAAV2-hIL-6 (hIL-6). The red nucleus was visualized by BDA (green) injection into specific coordinates 2 weeks before tissue harvest. Phospho-STAT3 is present in AAV2-hIL-6 but not AAV2-GFP treated mice. Scale bar: 50 µm

To test whether cortical motoneurons were synaptically connected with raphe neurons, we used AAV1, which can transsynaptically transduce supraspinal target neurons in rats^32^. We injected AAV1-Cre into the left sensorimotor cortex of *Rosa-tdTomato* (RFP) reporter mice and isolated the brain stem tissue after 2 weeks. RFP-expressing (transduced) serotonergic neurons were detected in both sides of the nucleus raphe pallidus (NRPa) and the nucleus raphe magnus (NRM) (Fig. 6 E-L), indicating a synaptic connection to cortical neurons. Hence, unilateral cortical AAV2-hIL-6 application transneuronally activates regenerative signaling pathways in ipsi- and contralateral raphe neurons by the release of the protein.

## Discussion

The current study shows significant locomotor recovery in an adult mammal by a single, unilateral application of AAV2-hIL-6 into the sensorimotor cortex after a complete SCC. While pre-injury-induced PTEN^-/-^ in cortical neurons failed to facilitate functional recovery, postinjury applied AAV2-hIL-6 promoted longer CST axon growth than PTEN^-/-^ and, additionally, regeneration of serotonergic axons in the RpST, enabling significant locomotor recovery of both hindlimbs. Moreover, we provide first direct evidence that cortical motoneurons innervate raphe neurons, thus allowing the axonal transport and synaptic release of highly potent hIL-6 to stimulate the regeneration of serotonergic fibers in the brain stem. Thus, a transneuronal application of highly active molecules is a new and powerful approach to activate regenerative signaling, particularly in neurons located in brain regions that are challenging to access but relevant for functional recovery after spinal cord injury.

In contrast to incomplete, less severe hemisection- or contusion-injury models, permitting spontaneous functional recovery with BMS scores ≥ 6^11, 21, 23^, the SCC, just as in a complete transsection, eliminates all axonal connections between the proximal and distal portions of the spinal cord. This injury model is therefore highly resistant towards functional recovery and shows only some spasm-based active flexion movements of hindlimbs (BMS score of ≤ 2). Hence, most previous treatment strategies in this injury model did not reach significant effects beyond a reduction of dieback of CST axons^34–36^. Only PTEN^-/-^ in cortical motoneurons enabled regeneration of CST-axons and synapse formation with interneurons but still failed to restore locomotion^10, 11, 33, 37, 38^. We confirmed these findings and used the PTEN^-/-^ model as a reference to validate the effects of AAV2-hIL-6. Although cortical PTEN^-/-^ showed a robust effect at shorter distances suggesting a stronger initiation of axon regeneration in the CST, hIL-6 promoted axons to regenerate much further. Moreover, only AAV2-hIL-6 treatment significantly improved serotonergic axon growth in the RpST and, remarkably, enabled locomotion recovery of both hindlimbs. These effects were achieved in mice with different genetic backgrounds by a single, unilateral application of AAV2-hIL-6 after the SCC, making this gene therapeutic approach also a potential strategy to facilitate spinal cord repair in the clinic.

PTEN^-/-^ and hIL-6 differently activated regenerative pathways. While PTEN^-/-^ expectedly activated PI3K/AKT/mTOR, AAV2-hIL-6 only induced JAK/STAT3-signaling in transduced and adjacent cortical neurons (Fig. 1) and via the transneuronal route also in raphe neurons. It is therefore conceivable that the combinatorial activation of both signaling pathways by PTEN^-/-^ and hIL-6 in cortical motoneurons would also synergistically result in stronger CST-regeneration than each treatment by itself as previously shown in the optic nerve^16–18^. These synergistic effects were, however, limited to the regeneration of the CST due to the effect of PTEN^-/-^ being restricted to the virally transduced motoneurons in the cortex. Moreover, the finding that hIL-6 alone did not measurably affect PI3K/AKT/mTOR activity but showed even stronger effects than PTEN^-/-^ suggests that extensive activation of AKT/mTOR, which is associated with a cancerogenic risk, is not essential to achieve a reduction of axonal dieback or improvement of CST regeneration. AAV2-hIL-6 did not activate the MAPK/ERK pathway either leading to the conclusion that its beneficial effect was mediated by STAT3 phosphorylation as previously shown for RGC-regeneration^12, 16, 17, 39^. Future experiments need to address the question whether a specific knockout or inhibition of STAT3 in cortical neurons reduces the hIL-6 effects on CST-regeneration in the SCC model or whether the effects of hIL-6 on JAK/STAT3 activation and axon regeneration can be further increased in combination with a specific knockout of SOCS3, which normally limits the activity of JAK/STAT3-signaling^16, 17^.

Although AAV2-hIL-6 enabled longer CST-regeneration than PTEN^-/-^, our data suggest that the beneficial effect on functional recovery mostly depended on the improved regeneration of serotonergic fibers of the raphe nuclei. This is because i) the regeneration of CST axons by PTEN^-/-^ alone did not enable hindlimb recovery, ii) BL6 mice showed similar effects on RpST-regeneration and recovery as that seen in Ola-PTEN-floxed animals but with less CST regeneration after AAV2-hIL-6 application, and iii) selective elimination of serotonergic fibers by DHT^24, 27, 28^ abolished most of the recovered locomotion after AAV2-hIL-6 treatment without affecting AAV2-GFP treated control animals, thereby also verifying the specificity of the neurotoxin. Consistent with this, the critical role of RpST regeneration for locomotor recovery has been reported after less severe spinal cord injuries, which allow some endogenous sprouting of serotonergic axons^24, 40–43^. Although these data demonstrate that AAV2-hIL-6-mediated functional recovery depends on the regeneration of serotonergic fibers, we cannot exclude the possibility that hIL-6 might have also stimulated other nuclei in the brain, such as the red nuclei, which also receive collateral input from cortical motoneurons^31^. Consistently, intracortical AAV2-hIL-6 application also induced STAT3 activation in the red nucleus, verifying the transneuronal hIL-6 delivery. Whether the regeneration of rubrospinal tracts was also affected or contributed to the beneficial effects of AAV-hIL-6 is currently unknown. Moreover, improved regeneration of the CST on top of the RpST could have contributed to the beneficial effect of hIL-6, despite CST regeneration induced by PTEN^-/-^ alone being insufficient. This would explain the finding that the stronger CST regeneration in the AAV2-hIL-6/PTEN^-/-^ group compared to hIL-6 animals correlated with an improvement seen in the BMS subscore, although the RpST regeneration remained similar. So improved CST-regeneration might only affect functional recovery on top of RpST regeneration, which first enables the basic walking behavior. Accordingly, CST axon regeneration reportedly improves voluntary movements and skilled locomotion in less severe pyramidotomy- and contusion-injury models, which leave other spinal tracts still intact^26, 44–46^. Future experiments need to investigate to what extent regeneration of serotonergic fibers alone can mimic the full AAV2-hIL-6 effect on functional recovery and whether it also reveals beneficial effects in less severe spinal cord injury models (e.g., pyramidotomy, hemisection, and contusion).

Although the collateral projection of cortical motor neurons into various brain areas, e.g., in the striatum and the thalamus^47–49^, as well as the red nucleus and reticular formation, are well documented^50–52^, the innervation of raphe nuclei in the brain stem has not yet been directly shown. Only data from a recent study constructing a connectome map of the whole brain point to a cortical projection of layer V motoneurons into the nucleus raphe magnus without clearly addressing this possibility^31^. The current study used AAV1, which can trans-synaptically transduce neurons^32^ and demonstrates in transgenic mice that cortical neurons project almost equally into the ipsi- and contralateral sides of the raphe nuclei. Finally, the activation of STAT3 (pSTAT3) in serotonergic raphe neurons by unilateral AAV2-hIL-6 application in the motor cortex in our study provides further evidence for this innervation target.

The current study also demonstrates that virally expressed hIL-6 is not only released from the soma to stimulate adjacent motoneurons in a paracrine fashion but that it is also transported over long distances in axons of either RGCs in the visual system or cortical neurons. The release of active hIL-6 at axonal terminals/synapses was verified in cell culture experiments and by pSTAT3 staining in visual target areas as well as directly in the raphe nuclei themselves. In this context, it is worth mentioning that hIL-6 is highly potent, so that even the smallest quantities can activate all types of neurons expressing gp130, the receptor to which it directly binds^18^.

The high potency of hIL-6 and the almost equal activation of neurons in the ipsi- and contralateral sides of the raphe nuclei also explains why the unilateral application improved the recovery of both hindlimbs to a similar degree, and why the bilateral application had no additional beneficial effect on RpST regeneration or functional recovery.

In conclusion, the finding that transneuronal application of hIL-6 enables functional recovery opens many new possibilities to further improve the functional outcome by combining it with other strategies, such as neutralizing extracellular inhibitors at the lesion site^8, 42, 53^ or bridging the lesion site with permissive grafts^54, 55^. These combinatorial strategies, also in less severe injury models, may lead to promising new methods for maximizing axon regeneration and functional recovery after spinal cord injury, potentially also in humans.

## Methods

### Mouse strains

Male and female mice were used for all experiments. Wild-type C57BL/6 and 129/Ola mice were cross-bred with PTEN^f/f^ mice (C57BL/6;129) to obtain C57BL/6;129/J-TgH (Pten-flox) animals. C57BL/6J mice were obtained from Janvier Labs. Rosa26 loxP-stop-loxP-tdTomato (*Rosa-tdTomato)* mice were obtained from Jackson Laboratories (Stock No: 007914). All animals were housed under the same conditions for at least ten days before the start of experiments and generally maintained on a 12 h light/dark cycle with *ad libitum* access to food and water. All experimental procedures were approved by the local animal care committee and conducted in compliance with federal and state guidelines for animal experiments.

### Intracortical injection of pups

Postnatal (P1) PTEN floxed (C57BL/6;129/J-TgH(Pten-flox)) pups were fixed under a stereotactic frame and continuously supplied with 2% isoflurane for anesthesia via a mouthpiece. A midline incision into the skin was made to expose the skull using microscissors. Since the skull of P1 mice is still soft, the cortex could be accessed using a 30-gauge needle to create two small holes in the left skull hemisphere with the following coordinates: −0.2 mm and 0.3 mm anteroposterior, 1.0 mm lateral to bregma. For AAV2-GFP or AAV2-Cre application, 770 nl of the virus suspensions were injected at a depth of 0.5 mm into the two sites using a pulled glass pipette connected to a nanoliter injector (Nanoject II, Drummond). To inject 770 nl, we applied 11 pulses of 70 nl at a rate of 23 nl/s and waited for 10 s after each pulse to allow distribution of the virus solution. After injection, the pipette was left in place for 1 minute before being carefully withdrawn, and the incision carefully closed with a 4-0 black silk suture.

### Intracortical injection of adult mice

For intracortical injection, adult PTEN floxed (C57BL/6;129/J-TgH(Pten-flox)), or adult wt (C57BL/6) mice were anesthetized by intraperitoneal injections of ketamine (120 mg/kg) and xylazine (16 mg/kg), and then placed in a stereotaxic frame. A midline incision was made over the skull to open the skin and to reveal the bregma. A microdrill with a 0.5 mm bit was used to open a 2 × 2 mm window on each side of the skull to expose the sensorimotor cortex. The respective AAV2 was injected into the cortex layer V through a glass pipette attached to a nanoliter injector (Nanoject II, Drummond). To this end, four injections of 490 nl each were given either unilaterally (left hemisphere) or in both hemispheres, at the following coordinates: 1.5 mm lateral, 0.6 mm deep, and 0.5 mm anterior; 0.0 mm, 0.5 mm, and 1.0 mm caudal to bregma. For injecting 490 nl into each injection site, we applied 7 pulses of 70 nl at a rate of 23 nl/s and waited for 10 s after each pulse to allow distribution of the virus solution. The needle was left in place for 1 minute before moving to the next site, and the brain was kept moist during the procedure by moving the skin over the exposed area after each injection. After surgery, the skin was closed with sutures. The virus transduced mainly neurons in layer 5 of the primary motor cortex (M1). We observed only very rare transduction of astrocytes or neurons of other M1 layers (Fig. S1 A-E). All GFP-expressing neurons also expressed hIL-6 (Fig. S1 F). Therefore, hIL-6 transduction was identified by GFP expression during the whole study.

### Injection into the red nucleus

Adult wt mice were anesthetized by intraperitoneal injections of ketamine (120 mg/kg) and xylazine (16 mg/kg) and then placed in a stereotaxic frame. A midline incision was made over the skull to open the skin and to reveal the bregma. A microdrill with a 0.5 mm bit was used to open a 1 × 1 mm window in the skull to expose the cortex. Biotinylated dextran amine (BDA, 10,000 MW, 10% solution in water, Invitrogen, D1956) was injected into the red nucleus through a glass pipette attached to a nanoliter injector (Nanoject II, Drummond). To this end 500 nl was given as previously described^26^ at the following coordinates: 0.6 mm lateral, 3.5 mm deep, and 2.5 mm caudal to bregma. For injecting 500 nl, we applied 5 pulses of 100 nl at a rate of 5 nl/s and waited for 10 s after each pulse to allow distribution of the virus solution. The needle was left in place for 1 minute before removing, and the brain was kept moist during the procedure. After surgery, the skin was closed with sutures.

### Complete spinal cord crush

For complete spinal cord crush, adult PTEN floxed (C57BL/6;129/J-TgH(Pten-flox)), or wt (C57BL/6) mice were anesthetized by intraperitoneal injections of ketamine (120 mg/kg) and xylazine (16 mg/kg). A midline incision of ∼1.5 cm was performed over the thoracic vertebrae. The fat and muscle tissue were cleared from thoracic vertebrae 7 and 8 (T7, T8). While holding onto T7 with forceps, we performed a laminectomy at T8 in order to expose the spinal cord. Afterward, the complete spinal cord was crushed for 2 s with forceps that had been filed to a width of 0.1 mm for the last 5 mm of the tips to generate a homogeneously thin lesion site. To ensure that the full width of the spinal cord was included we took care to gently scrape the forceps’ tips across the bone on the ventral side of the vertebral canal so as not to spare any axons ventrally or laterally. After surgery, the muscle layers were sutured with 6.0 resorbable sutures and the skin secured with wound clips. The completeness of the injury in the SCC model was verified by an astrocyte free gap and the absence of any spared CST or raphe spinal tract (RpST) fibers caudal to the lesion site shortly after injury (Fig. S7 E-J).

### CST tracing

To trace axons from cortical motoneurons of PTEN floxed (C57BL/6;129/J-TgH(Pten-flox)), or wt (C57BL/6) mice, we injected the axon tracer biotinylated dextran amine (BDA, 10,000 MW, 10% solution in water, Invitrogen, D1956) into the sensorimotor cortex 2 weeks before the mice were sacrificed. Therefore, the skin was opened, and 490 nl BDA were applied to each injection site using the same procedure and coordinates as described for AAV2 injection. After surgery, the skin was closed with sutures. In groups with the PTEN knockout, we verified that 70-80% of BDA-traced neurons were also Cre-positive (Fig. S7 A-D).

### DHT injection

Mice received bilateral intracerebroventricular injections of 30 µg of the serotonin neurotoxin 5,7-dihydroxytryptamine (DHT, Biomol) dissolved in 0.5 μl of 0.2% ascorbic acid in saline to deplete serotonergic inputs to the lumbar spinal cord. To this end, mice were anesthetized and fixed under a stereotactic frame as described above for intracortical injections. The tip of a glass micropipette attached to a nanoliter injector (nanoject II, Drummond) was positioned at the following coordinates: 0.6 mm posterior, 1.6 mm lateral to bregma, and 2 mm deep from the cortical surface. DHT was injected at the same rate as described for the AAV2 injections, and the pipette was left in place for 1 minute before the withdrawal. Thirty minutes before DHT injection, the monoamine uptake inhibitor, desipramine (Sigma), was administered at 25 mg/kg intraperitoneally to prevent the uptake of DHT into noradrenergic neurons.

### Intraocular injection of AAV2

Adult mice (C57BL/6) received intravitreal AAV2 injections (1 µl) as described previously^18, 56^. After three weeks, axons of the visual pathway were labeled by intravitreal injection of 2 µl of Alexa Fluor 555-conjugated cholera toxin β subunit (0.5% CTB, in PBS; Molecular Probes, Carlsbad, Eugene, USA) and tissues isolated after another week.

### AAV virus production

AAV1-Cre virus was obtained from Addgene (pAAV.CMV.HI.eGFP-Cre.WPRE.SV40; plasmid #105545). AAV2 viruses were produced in our laboratory. To this end, AAV plasmids carrying either cDNA for Cre-HA, GFP, or hIL-6 downstream of a CMV promoter were co-transfected with pAAV-RC (Stratagene) encoding the AAV genes rep and cap, and the helper plasmid (Stratagene) encoding E24, E4, and VA into AAV-293 cells (Stratagene) for recombinant AAV2 generation. Purification of virus particles was performed as described previously^12^. Mainly, cortical neurons are transduced upon intracortical injection of AAV2 into layer V, as this virus serotype is highly neurotropic.

### Tissue processing

Mice were anesthetized and perfused through the heart with cold PBS followed by 4% PFA in PBS. Isolated brains and spinal cords were post-fixed in 4% PFA overnight, transferred to 30% sucrose at 4°C for five days. The bulk of the brain, medulla, and different segments of the spinal cord were embedded in Tissue-Tek (Sakura) and frozen at −20°C. For spinal cords subjected to T8 crush, a ∼1.2 cm segment from 3 mm rostral to- and 9 mm caudal from the injury was embedded for sagittal sections. Spinal cord tissues of ∼2 mm length just rostral to-and caudal from this 1.2 cm segment were embedded for transverse sections to evaluate axon sprouting proximal to the lesion site (rostral segments), and completeness of the lesion by identification of potentially spared axons (caudal segments). Tissues were sectioned on a cryostat (Leica) at a thickness of 20 μm, thaw-mounted onto Superfrost plus slides (ThermoFisher), and stored at −20°C until further use. Transverse sections of the medullary pyramids ∼1 mm above the pyramidal decussation were cut at a thickness of 14 µm to quantify the total number of BDA traced corticospinal axons (see below).

### DRG neuron two-compartment cultures

Dorsal root ganglion-(DRG) neurons were harvested from adult mice, as previously described^57^. In brief, isolated DRG (T8-L5) were incubated in 0.25% trypsin/EDTA (GE Healthcare) and 0.3% collagenase Type IA (Sigma) dissolved in DMEM (Invitrogen) at 37°C and 5% CO_2_ for 45 min and mechanically dissociated afterward. Cells were resuspended in DMEM containing B27-supplement (Invitrogen 1 : 50) and penicillin/streptomycin (500 U/ml; Merck, Millipore) and seeded into the somal compartment of microfluidic two-compartment chambers (AXIS Axon Isolation Device, Millipore), mounted on poly-D-lysine (0.1 mg/ml, molecular weight 70,000-150,000 kDa; Sigma) plus laminin-coated (20 μg/ml; Sigma) culture dishes according to the manufacturer’s instructions. Neurons were cultured at 37°C and 5% CO_2_ for 3 days to allow axon extension through the microchannels and then transduced with hIL-6 expressing baculoviruses (BV) or GFP expressing control BV. Diffusion of particles or proteins through the microchannels into the axonal compartment was prevented by using a hydrostatic pressure due to two different volumes of medium with a resulting antagonistic microflow of liquid between the axonal and the somal chambers. After 24 h, the virus-containing medium was replaced with fresh culture medium (DMEM, 1:50 B27 supplement, and 1:50 penicillin/streptomycin). After 48 hours the medium of 5 axonal compartments from either BV-GFP or BV-hIL-6 transduced cells were collected respectively and concentrated by 10-fold, using Merck Amicon centrifugal filter units with a retaining size of 30 kDa. The concentrated medium was then used to culture retinal ganglion cells (RGCs), as described in the following paragraph. After fixation with 4% PFA for 30 min at RT, cell cultures were processed for immunocytochemical staining with antibodies against synapsin (1:1000, Millipore, RRID:AB_2200400), GFP (1:500; Novus; RRID:AB_10128178), and IL-6 (1:500; Abcam; RRID:AB_2127460). Stained axons were photographed using a confocal microscope (630 x, SP8, Leica)

### Dissociated retinal cell cultures

To prepare primary retinal cell cultures, tissue culture plates (4-well plates; Nunc, Wiesbaden, Germany) were coated with poly-D-lysine (PDL, 0.1 µg/ml, molecular weight between 70,000 and 150,000 Da; Sigma, St Louis, USA) and with 20 mg/ml laminin (Sigma). To prepare low-density retinal cell cultures, mice were killed by cervical dislocation. Retinae were rapidly dissected from the eyecups and incubated at 37 °C for 30 min in a digestion solution containing papain (10 U/mL; Worthington) and L-cysteine (0.2 μg/mL; Sigma) in Dulbecco’s modified Eagle medium (DMEM; Thermo Fisher). Retinae were then rinsed with DMEM and triturated in 1.5 ml medium obtained from axonal compartments of DRG neuron cultures as described above. Dissociated cells were passed through a cell strainer (40 µm, BD Falcon, Franklin Lakes, USA), and 300 µl of cell suspension was added into each well of the coated culture plates. Retinal cells were cultured for 48 h and then fixed with 4% PFA (Sigma).

After fixation with 4% PFA for 30 min at RT, cell cultures were processed for immunocytochemical staining with a βIII-tubulin-antibody (1:2,000, BioLegend, RRID:AB_2313773). All RGCs with regenerated neurites were photographed using a fluorescent microscope (200 ×, Axio Observer.D1, Zeiss), and neurite length was determined using ImageJ software. Also, the total number of βIII-tubulin-positive RGCs with an intact nucleus (6-diamidino-2-phenylindole (DAPI)) per well was quantified to test for potential neurotoxic effects. The average neurite length per RGC was determined by dividing the sum of neurite length per well by the total number of RGCs per well. Cultures were arranged in a pseudo-randomized manner on the plates so that the investigator would not be aware of their identity. Data represent means ± SEM. of two independent experiments, each with four wells as technical replicate per treatment.

### Immunohistochemistry

Cryosections of brain, medulla and spinal cord were thawed, washed in PBS for 15 minutes and permeabilized by 10 minutes incubation in 100% Methanol (Sigma). Sections were stained with antibodies against the HA-tag (1:500; Sigma; RRID:AB_260070), pSTAT3 (1:200; RRID: AB_2491009), pAKT-Thr308 (1:500; RRID: AB_2629447), pERK1/2 (1:500; RRID: AB_331646), pS6 (1:500; RRID: AB_2181035) (all Cell Signaling Technology), GFP (1:500 Thermo Fischer, RRID: AB_221570), GFP (1:500; Novus; RRID:AB_10128178), IL-6 (1:500; Abcam; RRID:AB_2127460) GFAP (1:50; Santa Cruz Biotechnology; RRID: AB_627673), serotonin (1:5,000; Immunostar; RRID: AB_572263, RRID:AB_572262) and NeuN (1:2,000; Abcam, AB_10711040). Secondary antibodies included anti-mouse, anti-goat and anti-rabbit conjugated to Alexa Fluor 405 (1:500; Jackson ImmunoResearch), 488, or 594 (1:1,000; Invitrogen), respectively. BDA traced CST neurons and axons were detected using streptavidin fluorescently labeled with Alexa Flour 405, 488, or 594 (1:500, Thermo Fischer). Sections were embedded in Mowiol (Sigma) and analyzed using fluorescent widefield (Observer.D1, Zeiss; Axio Scan.Z1, Zeiss), or confocal laser scanning (SP8, Leica) microscopes.

### Western blot

For cortical lysate preparation, mice were killed, and a cuboid piece of cortical tissue of 1.5 × 2 mm with 1 mm depth from the cortical surface was dissected and isolated from the sensorimotor cortex around the coordinates used for viral injection. For lysates of brain stem raphe nuclei, a 2.5 mm long and 1.5 mm wide piece of tissue was isolated around the midline of the medulla with a depth of 1 mm starting above pyramidal tracts, which were thereby also removed. Tissues were homogenized in lysis buffer (20 mM Tris-HCl, pH 7.5, 10 mM KCl, 250 mM sucrose, 10 mM NaF, 1 mM DTT, 0.1 mM Na_3_VO_4_, 1% Triton X-100, 0.1% SDS) with protease inhibitors (Calbiochem) and phosphatase inhibitors (Roche) using five sonication pulses at 40% power (Bandelin Sonoplus). Lysates were cleared by centrifugation in a tabletop centrifuge (Eppendorf) at 4,150 × g for 10 minutes at 4 °C. Proteins were separated by SDS/PAGE, using Mini TGX gels (10%, Bio-Rad) according to standard protocols, and then transferred to nitrocellulose membranes (0.2 µm; Bio-Rad). Blots were blocked in 5% dried milk in Tris- or phosphate-buffered saline solution with 0.05% Tween-20 (TBS-T and PBS-T, respectively; Sigma) and incubated with antibodies against pSTAT3 (1:200; RRID: AB_2491009), IL-6 (1:1000; Abcam; RRID:AB_2127460), pAKT-Thr308 (1:500; RRID: AB_2629447), pERK1/2 (1:500; RRID: AB_331646), pS6 (1:500; RRID: AB_2181035) (all Cell Signaling Technology), GFP (1:500 Thermo Fischer, RRID: AB_221570), or against β-actin (1:5000; AC-15; Sigma, RRID: AB_476744) at 4 °C overnight. All primary antibodies were diluted in TBS-T or PBS-T containing 5% BSA (Sigma). Anti-rabbit or anti-mouse IgG conjugated to HRP (1:80,000; Sigma) were used as secondary antibodies. Primary antibodies against pSTAT3 and pAKT were detected by using a conformation-specific HRP-conjugated anti-rabbit IgG secondary antibody (1:4,000; Cell Signaling Technologies; RRID: AB_10892860). Antigen-antibody complexes were visualized by using enhanced chemiluminescence substrate (Bio-Rad) on a FluorChem E detection system (ProteinSimple). Western blots were repeated at least three times to verify results. Band intensities were quantified relative to respective loading controls by using ImageJ software.

### Tissue clearing

Brain and spinal cord tissue were isolated from mice after perfusion with PBS and 4% PFA as described above. Tissue was postfixed overnight at 4 °C before further processing. Wholemount immunostaining was performed as described previously^58^. In brief, tissue was washed two times for 1 h in PBS containing 0.2% Triton-X-100 and then incubated overnight at 37 °C in PBS containing 0.2% TritonX-100 and 20% DMSO. Afterwards, the tissue was again incubated overnight at 37°C in PBS containing 0.1% Tween-20, 0.1% Triton-X-100, 0.1% deoxycholate, 0.1% NP40 and 20% DMSO. After two washing steps in PBS with 0.2% TritonX-100, specimens were permeabilized (PBS + 0.2% TritonX-100 + 23% glycine + 20% DMSO) for 1 day at 37°C, blocked (PBS + 0.2% TritonX-100 + 6% donkey serum + 10% DMSO) at 37°C for 1 d, and incubated with primary antibodies against pSTAT3 (1:100; Cell Signaling Technology; RRID: AB_2491009, RRID:AB_2198588), GFP (1:500; Thermo Fischer, RRID: AB_221570), serotonin (1:2,500; Immunostar; RRID: AB_572263, RRID: AB_572262) and NeuN (1:500; Abcam, RRID: AB_10711040) at 37°C in blocking solution for 2 d. Afterward, tissue was incubated for another 2 d at 37°C with fluorescently labeled streptavidin (Alexa Flour 405, 488, or 594; 1:250; Thermo Fischer) to detect BDA traced CST axons, or with secondary antibodies including anti-mouse, anti-goat and anti-rabbit conjugated to Alexa Fluor 405 (1:250; Jackson ImmunoResearch), 488 or 594 (1:500; Invitrogen), respectively. Subsequently, spinal cords and brains were dehydrated in ascending concentrations of tetrahydrofuran (50%, 80%, 100%), incubated in dichloromethane and then transferred to a clearing solution (benzyl alcohol: benzyl benzoate (1:2)) and imaged with a confocal laser scanning microscope (SP8, Leica).

### Quantification of the total number of BDA traced CST axons

To obtain the number of BDA traced axons, four adjacent medullary cross-sections, ∼1 mm rostral to the pyramidal decussation, were stained with fluorescently labeled streptavidin and imaged with a fluorescent microscope. Four 25 × 25 μm squares were randomly superimposed on each image, and the number of labeled axons in every square was counted and averaged. The area of the entire pyramid and the total area inside a counted square were measured and used to extrapolate the total number of BDA-labeled axons per pyramid.

### CST sprouting proximal to the lesion site

The sprouting index was determined as described previously^10, 45^. In brief, images of transverse spinal cord sections ∼3 mm rostral to the lesion site from untreated controls and mice eight weeks after spinal cord crush were taken. To quantify the number of sprouting axons, we drew a horizontal line through the central canal and across the lateral rim of the gray matter. Three vertical lines were then drawn to divide the horizontal line into three equal parts (Z1, Z2, Z3), starting from the central canal to the lateral rim. Only fibers crossing these lines were counted in each section. The results were presented after being normalized against the number of total CST fibers at the medulla level obtained as described above. At least three sections were quantified for each mouse.

### Evaluation of CST-axon retraction

The retraction of axons rostral to the injury site was quantified as an axon density index by measuring the BDA staining intensity as described previously^10, 11, 33^. A series of 100 μm wide rectangles covering the entire dorsoventral axis of the spinal cord were superimposed onto sagittal sections, starting from 1.5 mm rostral up until the injury site, which was visible by its morphology and additional GFAP staining of the glial scar. The background signal in BDA negative areas was measured in each section and subtracted from quantified BDA staining. Additionally, the pixel intensity values of each rectangle were normalized against the intensity at 1.5 mm rostral to the lesion site. The Axon Density Index was then plotted as a function of the distance to the injury. All sagittal sections containing the main CST from each animal were quantified and averaged.

### Evaluation of CST regeneration

Sagittal sections of spinal cord segments starting from 3 mm caudal to and up until 9 mm distal to the lesion site were stained for BDA using fluorescently labeled streptavidin as described above. Although the injury site was visible by morphology, at least one section per animal was stained against GFAP to ensure the identity of the lesion edges. Axon regeneration was quantified from stitched images scanned under a 10× objective on a fluorescent Axio Scan.Z1 microscope (Zeiss). A grid with lines spaced every 100 µm was aligned to each image, and the number of axons at indicated distances from the caudal edge of the lesion were quantified in ∼40 adjacent sections per animal. For quantification, all axons observed within an area of 0.5 mm caudal and rostral to the respective measuring point were counted. To obtain the axon index and to control for variations in tracing efficiency, the total number of axons at each distance was divided by the total number of BDA-labeled axons in the brain stem. Only animals with a complete lesion, proven by the absence of any BDA stained axons in spinal cord cross-sections ∼10-11 mm distal to the lesion site, where included in the evaluation.

### Evaluation of RpST fiber regeneration

Sagittal spinal cord sections as described above were stained against serotonin and imaged under a fluorescent microscope (10x, Axio Scan.Z1, Zeiss). Images were superimposed with a grid as described for CST regeneration. Axons that reached indicated distances from the distal border of the lesion site were quantified in ∼40 sections (every second section) per animal, averaged over all analyzed sections and then extrapolated to the whole number of sections per spinal cord. For quantification, all axons observed within an area of 0.5 mm caudal and rostral to the respective measuring point were counted.

### Basso Mouse Scale (BMS)

The locomotory behavior of mice was tested and scored according to guidelines of the Basso Mouse Scale^21^. Therefore, each mouse was placed separately in a round open field of 1 m in diameter and observed by two testers for 4 minutes. Scoring was based on different parameters such as ankle movements, paw placement, stepping pattern, coordination, trunk instability, and tail position, with a minimum score of 0 (no movement) to a maximum score of 9 (normal locomotion). For animals that have attained frequent plantar stepping (BMS ≥ 5), we additionally determined the BMS subscore which discriminates more precisely the fine details of locomotion such as coordination or paw position that may not be differentiated by the BMS^11, 21^. The subscore starts with a minimum score of 0 whereas the maximum value is 11. Before testing, mice were acclimated to being handled and the open field environment. BMS tests were performed before the injury, on days 1, 3, 7, and then weekly (over eight weeks) after spinal cord injury. A BMS score of 0 at 1 d after injury and the absence of BDA staining in distal spinal cord cross-sections, as described above were used as quality criteria for completeness of the lesion. Mice that did not meet these criteria were excluded from the analysis.

### Statistics

Significances of intergroup differences were evaluated using Student’s t-test or analysis of variance (ANOVA) followed by Holm-Sidak, or Tukey post hoc test using the Sigma STAT 3.1 software (Systat Software). Statistical significance of intergroup differences was defined as the following: *p < 0.05; **p < 0.01; ***p < 0.001. Statistical details for individual experiments are presented in the corresponding figure legend.

## Supporting information

Video S1: Open field locomotion after SCC and AAV2-GFP treatment The video shows a PTEN-floxed Ola mouse (PTEN+/+) that received an injection of AAV2-

Video S2: Open field locomotion after SCC and AAV2-hIL-6 treatment The video shows a PTEN-floxed Ola mouse (PTEN+/+) that received an injection of AAV

Video S3: Open field locomotion after PTEN-/- and SCC The video shows a PTEN-floxed Ola mouse that received an injection of AAV2-Cre (PTEN-/-) into t

Video S4: Open field locomotion after PTEN-/- and SCC with AAV2-hIL-6 treatment The video shows a PTEN-floxed Ola mouse that received an injection of

Video S5: Open field locomotion after DHT- mediated depletion of raphe spinal input The video shows a PTEN-floxed Ola mouse (PTEN+/+) that was subjec

Video S6: Open field locomotion determined in a BL6 mouse after SCC and AAV2-hIL-6 treatment The video shows a C57BL/6J wildtype mouse (BL6) that was

Video S7: Intracortical hIL-6 application induces STAT3 phosphorylation in raphe nuclei Three-dimensional projection of transverse confocal scan throu

---

**Supplementary Figure 1:**
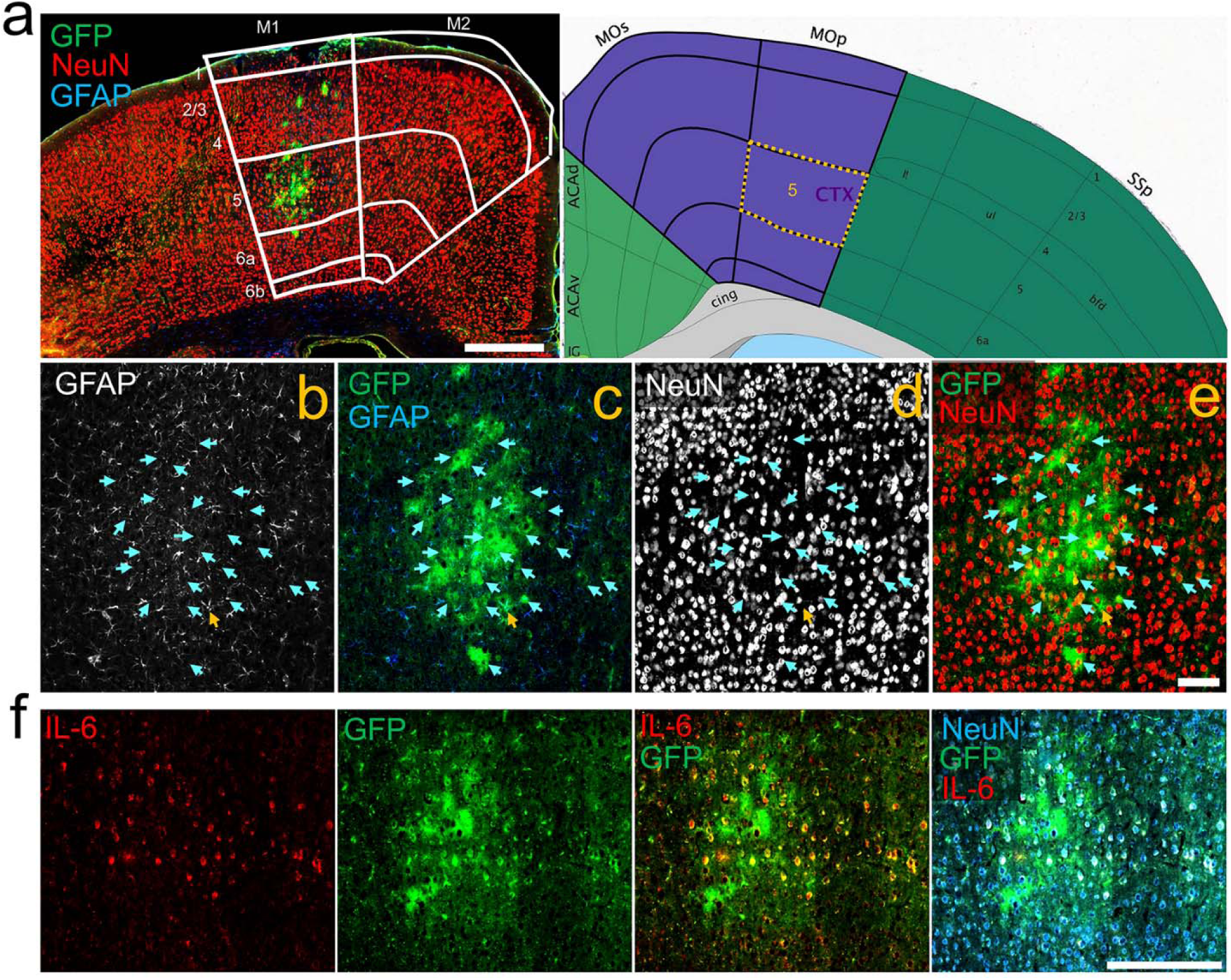
Validation of cortical AAV2 transduction. A) Coronal section of the sensorimotor cortex 3 weeks after AAV2-hIL-6 injection stained for glial fibrillary acid protein (GFAP, blue), NeuN (red), and GFP (green). White boxes indicate layers of primary (M1) and secondary (M2) motor cortex in an overlay with a corresponding map from *Allen Brain Atlas* (right). Scale bar 500 µm. B-E) Higher magnification of the image shown in A. Blue arrows mark NeuN positive neurons, while orange arrows indicate GFAP positive astrocytes. Scale bar: 50 µm. F) Coronal section of the sensorimotor cortex as described in A stained for hIL-6 (IL-6, red), GFP (green) and NeuN (blue). Scale bar: 200 µm

**Supplementary Figure 2:**
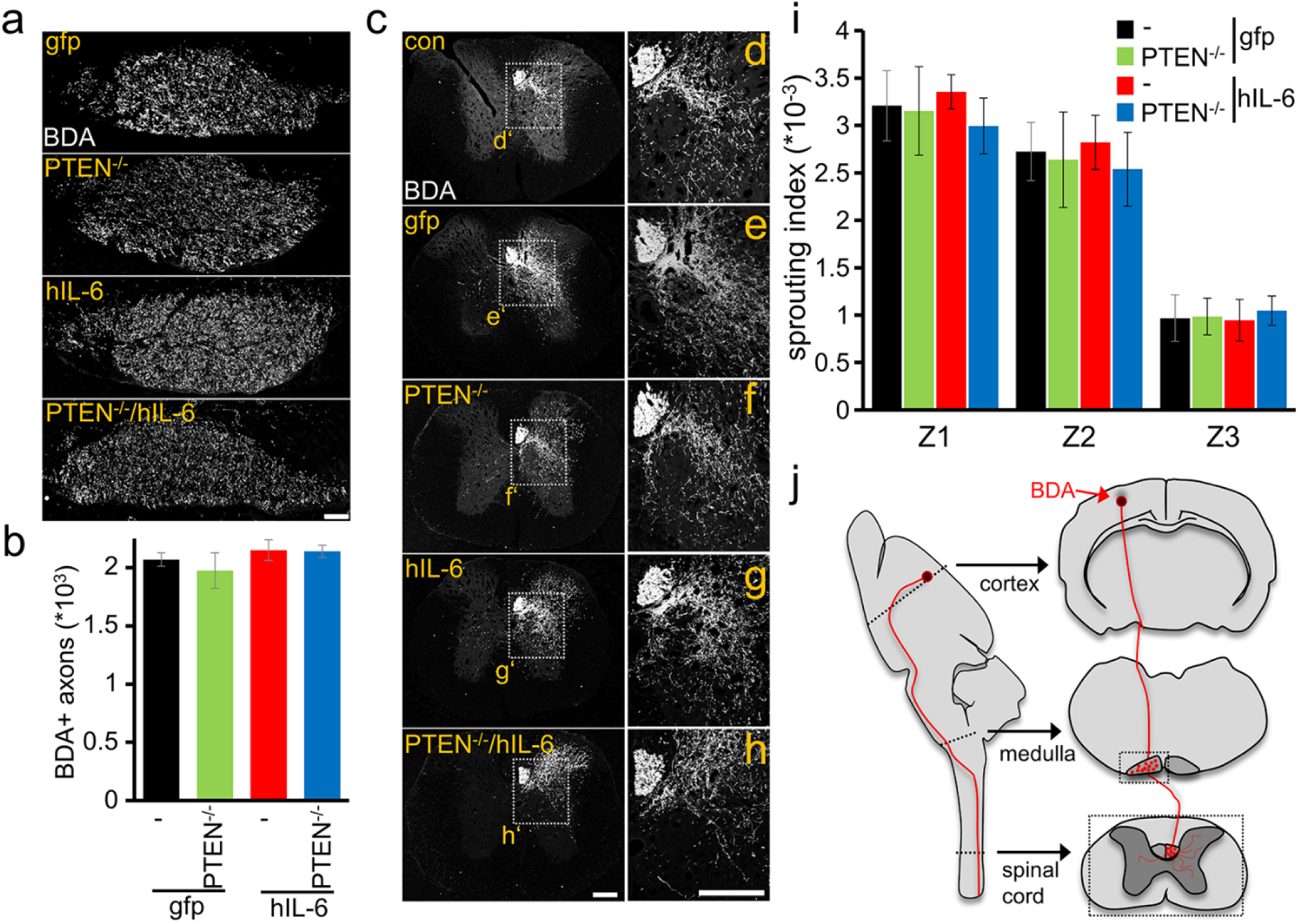
hIL-6 does not affect proximal CST axon sprouting. **A)** Coronal sections of the medullary pyramid with BDA-labeled CST axons in PTEN^f/f^ mice <u>8</u> weeks after spinal cord crush (<u>SCC</u>). Animals had received intracortical injections of either AAV-GFP (PTEN^+/+^) or AAV2-Cre (PTEN^-/-^) at P1. At the age of 8 weeks, animals were subjected to <u>SCC</u> and subsequently received intracortical AAV2-hIL-6 (hIL-6) or AAV2-GFP (<u>gfp</u>) injections. BDA tracing was performed 2 weeks before tissue isolation. Scale bar: 50 µm. **B)** Quantification of the density of BDA-labeled CST-axons in sections as described in **A**. Counts of five sections were averaged per animal. Values represent means +/- SEM of 5-9 animals per group (PTEN^+/+^/gfp, n=5; PTEN^+/+^/hIL-6, n=9; PTEN^-/-^/gfp, n=6; PTEN^-/-^/hIL-6, n=9). **C)** Representative images are showing BDA-labeled CST-axons in transverse spinal cord sections 3 mm rostral to the lesion site in animals as described in **A** and additional wt mice without SCC (con). Scale bar: 100 µm. **D-H)** Higher magnification of dashed boxes from images in **C**. Scale bar: 200 µm. **I)** Quantification of axons sprouting into the ipsilateral gray matter at defined distances from the midline (Z1-3). Counts of 5 sections were averaged per animal. Values represent means +/- SEM of 5-9 animals per group (PTEN^+/+^/gfp, n=5; PTEN^+/+^/hIL-6, n=9; PTEN^-/-^/gfp, n=6; PTEN^-/-^/hIL-6, n=9). **J)** Schematic drawing illustrating the location of coronal sections of the medulla and spinal cord shown in **A** and **C** (dotted boxes). Statistics in **B** and **I** were evaluated using two-way analysis of variance (ANOVA) with Holm-Sidak and Tukey post hoc tests, showing no significant differences between treatment groups.

**Supplementary Fig. 3:**
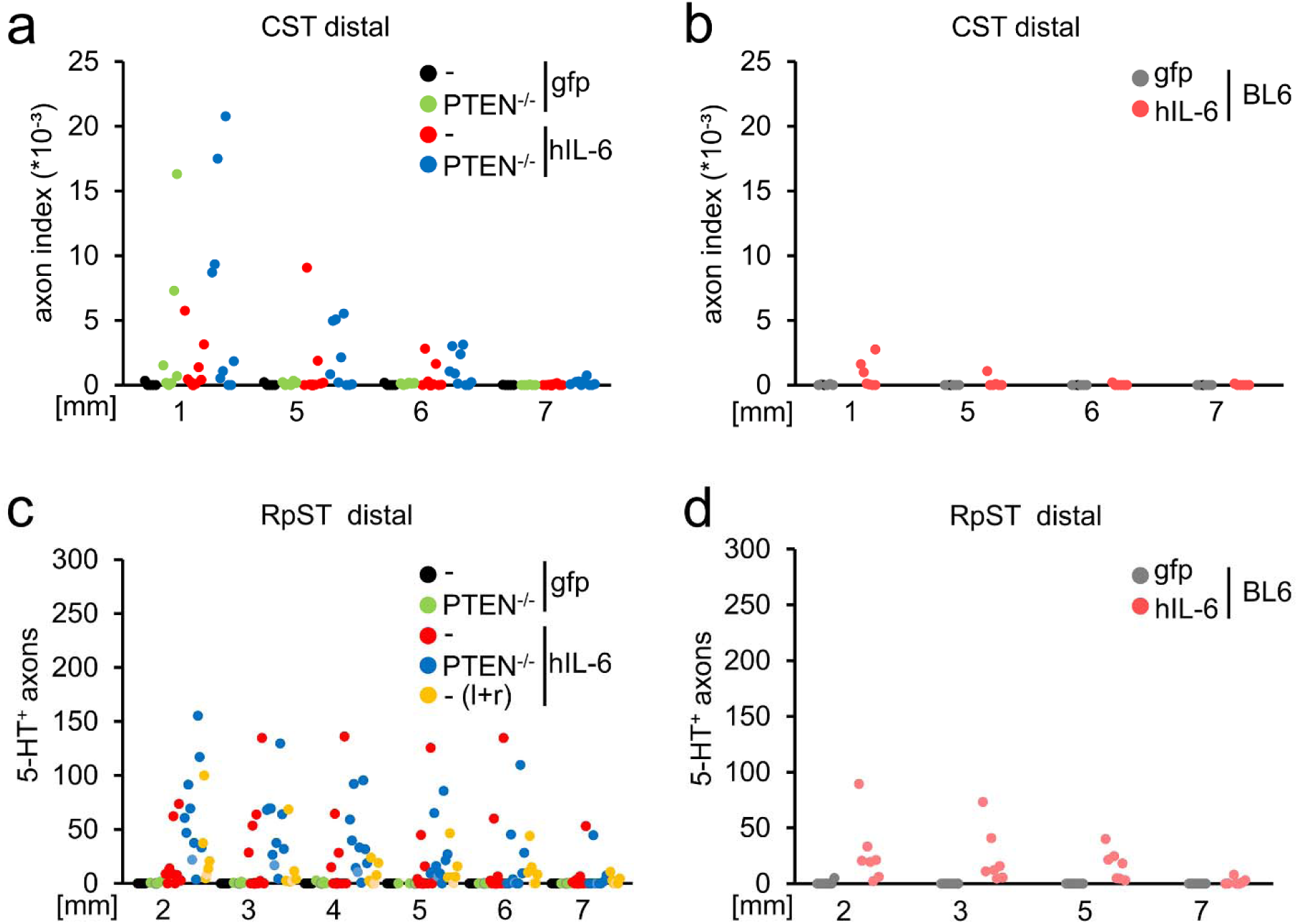
Values of individual animals in CST and RpST axon regeneration. **A/B)** Axon index of regenerating CST axons from individual animals with an Ola background as presented in Fig. 1 **I** (**A**) or BL6 animals, as presented in **Fig. S5 B** (**B**). **C/D)** Quantification of regenerating serotonergic RpST axons from single animals with Ola background as presented in Fig. 4 J (**C**), or BL6 animals as presented in **Fig. S5 C** (**D**).

**Supplementary Figure 4:**
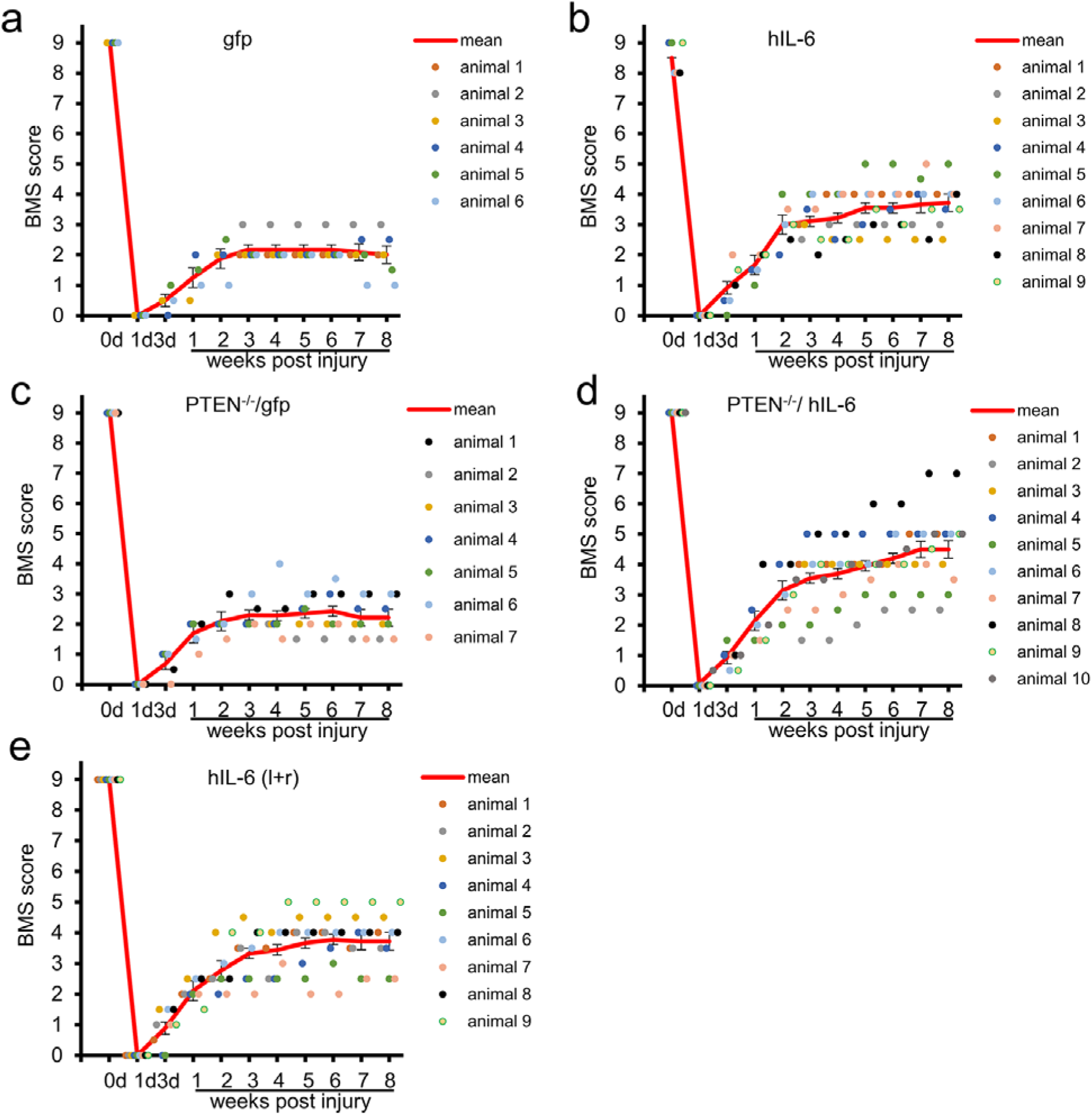
BMS Values of individual animals in open field analysis. **A-D)** Average BMS scores of left and right hind paws from individual animals treated as described in Fig. 3 **A** and their respective mean values (red line) as presented in Fig. 3 B. **E)** Average BMS scores of left and right hind paws and respective mean values (red line) from individual animals that received spinal cord crush and intracortical AAV2-hIL-6 injections in both hemispheres as presented in Fig. 3 F.

**Supplementary Figure 5:**
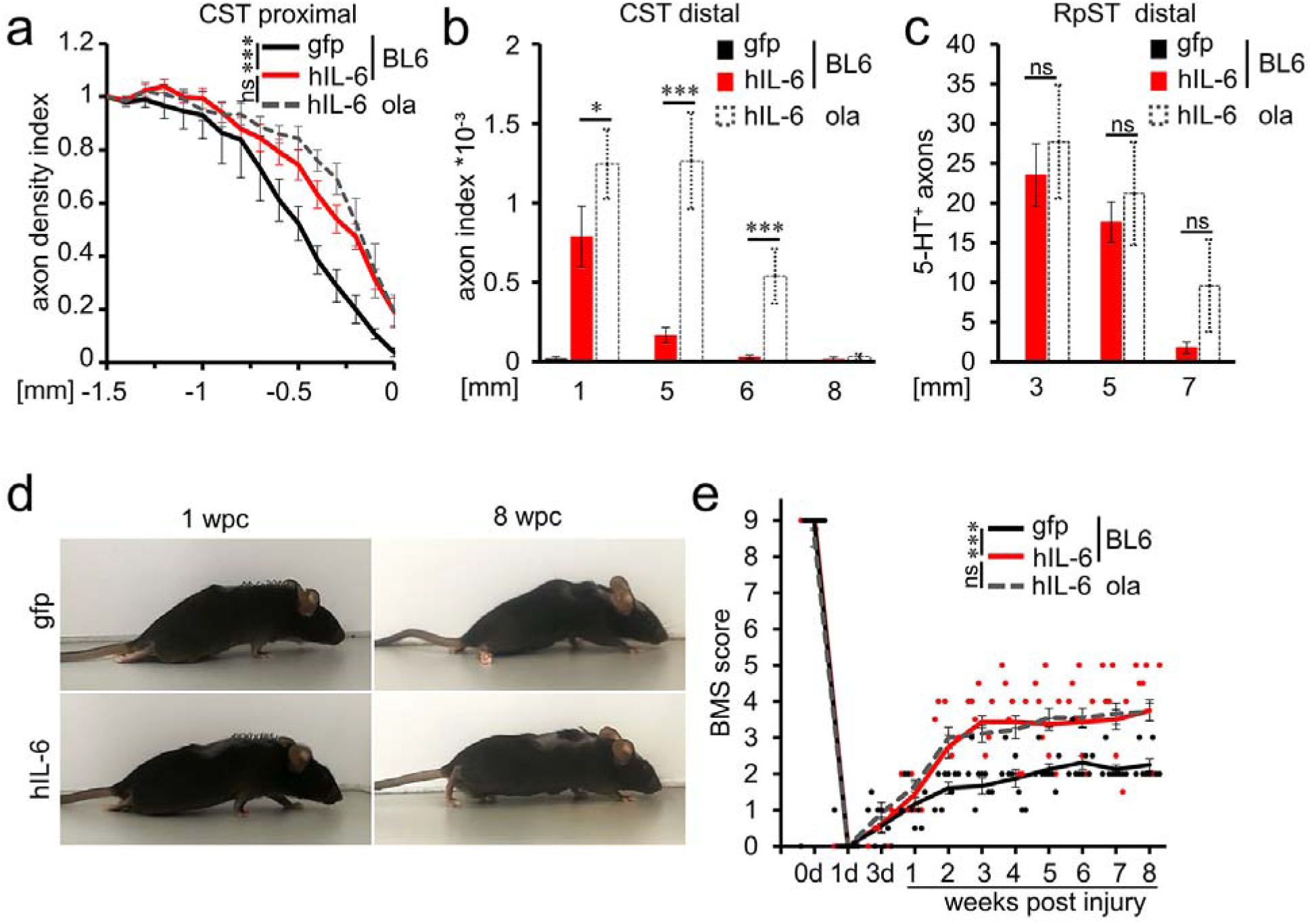
Hyper-IL-6 promotes functional recovery in different mouse strains. **A/B)** Quantification of BDA-traced CST fibers proximal- (**A**), and regenerated axons distal (**B**) to the lesion site of non-transgenic BL6 mice 8 weeks after spinal cord crush (wpc). Animals received injections of either AAV2-GFP (gfp) or AAV2-hIL-6 (hIL-6) into the left sensorimotor cortex immediately after SCC. The dashed line represents values of hIL-6 treated PTEN-floxed (PTEN^+/+^) Ola mice from Fig 1 for comparison. Values represent means +/- SEM of 7-8 animals per group (gfp, n=7; hIL-6, n=8) **C)** Regenerated 5-HT-positive RpST axons at indicated distances from the injury site in BL6 animals as described in **A**. Dashed bars represent values of hIL-6 treated PTEN-floxed (PTEN^+/+^) Ola mice from Fig. 4 J for comparison. Values represent means +/- SEM of 7-8 animals per group (gfp, n=7; hIL-6, n=8) **D)** Representative pictures showing open field movement of non-transgenic BL6 mice as described in **A** 1 and 8 weeks after SCC (wpc). **E)** BMS score of animals as described in **A** at indicated time points after SCC. Dots represent values of individual animals from the gfp (black) and the hIL-6 (red) group. The dashed gray line shows values of AAV2-hIL-6-treated PTEN-floxed (PTEN^+/+^) Ola animals from Fig. 3 B for comparison. Values represent means +/- SEM of 8-9 animals per group (gfp, n=8; hIL-6, n=9), showing the average score of the left and right hind paws. Significances of intergroup differences were evaluated at 8 weeks post-injury using a one-way analysis of variance (ANOVA) with Tukey or Holm-Sidak post hoc tests. Treatment effects as indicated: ***p<0.001; ns=non-significant.

**Supplementary Figure 6:**
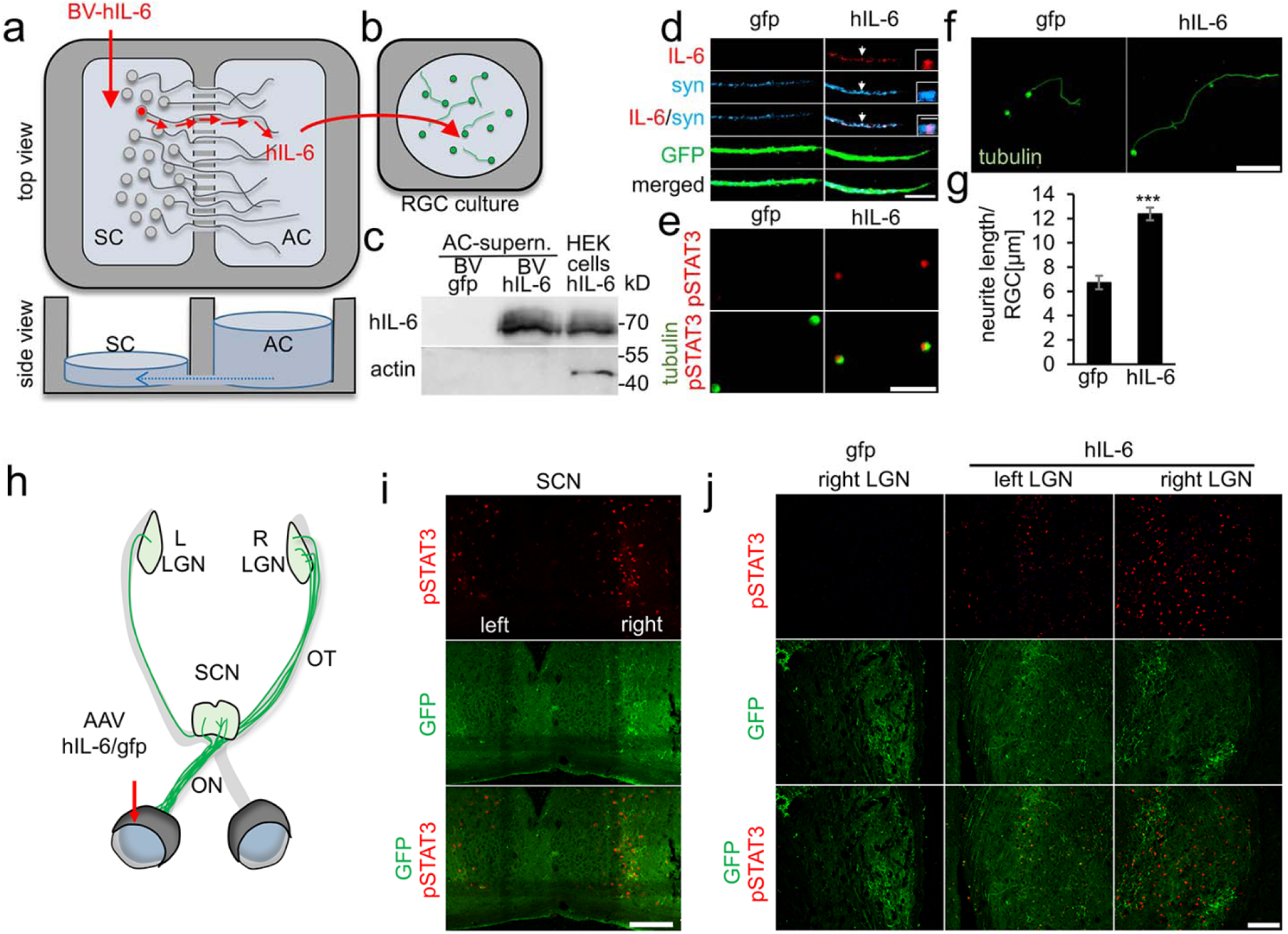
Axonal transport and release of virally expressed hIL-6. **A/B)** Schematic illustration of the experiment shown in **C-G**. Primary adult dorsal root ganglion-(DRG) neurons were cultured in microfluidic two-compartment chambers. Cell bodies in the soma compartment (SC) were transduced with a baculovirus expressing hIL-6 (BV-hIL-6) or with a GFP expressing control virus (BV-gfp). The diffusion of molecules from the SC to the axonal compartment (AC) was prevented by an antagonistic microflow (blue dotted arrow) due to different volumes of the medium in the AC and SC. Hyper-IL-6 is transported within axons of transduced neurons and released at axon terminals in the AC **(A)**. After 48 hours, media from AC of BV-gfp or BV-hIL-6 treated cultures were collected and used for culturing primary retinal ganglion cells (RGCs) **(B)**. **C)** Western blot analysis of the supernatant from the axonal compartment (AC-supern.) of cell cultures as described in A, showing the presence of hIL-6 after BV-hIL-6 but not BV-gfp treatment. Lysates of HEK293 cells transduced with BV-hIL-6 were used as a positive control. The lack of β-actin in the AC-supernatant proves the absence of cell fragments in the lysate, indicating the release of hIL-6 into the medium. **D)** Representative images of GFP (green) stained axonal tips from BV-gfp (gfp) or BV-hIL-6 (hIL-6) treated cultured DRG neurons as described in **A** stained for hIL-6 (IL-6, red) and synapsin (blue). Hyper-IL-6-positive vesicles were detected in only in axons of BV-hIL-6 treated neurons. White arrow indicates area magnified in insets. Scale bar: 10 µm, insets: 2µm. **E)** Representative images showing phospho-STAT3 (pSTAT3, red) in nuclei of βIII-tubulin (tubulin, green) positive RGCs 1 day after culturing in medium derived from BV-hIL-6 (hIL-6) treated DRG neuron cultures as described in **A, B**. Medium from BV-gfp (gfp) revealed no STAT3 activation. Scale bar: 50 µm. **F)** Representative images of neurite growth in βIII-tubulin (tubulin, green) stained RGCs as described in **A/B/D** exposed to hIL-6 or gfp-conditioned media. Scale bar: 100 µm. **G)** Quantification of neurite length per RGC in retinal cultures as depicted in E. Data represent means ± SEM of two independent experiments, each with four wells as technical replicate per treatment (n=8). Significances of intergroup differences were evaluated using Student’s t-test. Treatment effects: ***p<0.001 **H)** Illustration of the mouse visual pathway with the optic nerves (ON), optic tracts (OT), the suprachiasmatic nucleus (SCN) and lateral geniculate nuclei (LGN). 95% of transduced RGCs (green) of the left eye project axons into the right LGN; only 5% into the left hemisphere. Intravitreal AAV2-hIL-6 injection is indicated by the red arrow. **I)** Coronal section of both hemispheres of the SCN immunohistochemically stained for phosphorylated STAT3 (pSTAT3, red) from mice 3 weeks after intravitreal AAV2-hIL-6 injection. Staining was detected near GFP-positive axons of AAV2-hIL-6 transduced RGCs and was predominantly in the right hemisphere. Scale bar: 100 µm. **J)** Coronal sections of the left and right LGN 3 weeks after intravitreal injection of either AAV2-hIL-6 (hIL-6) or AAV2-GFP (gfp) into the left eyes. Phospho-STAT3 (pSTAT3, red) was only detected in hIL-6-treated animals near GFP-positive axons (green) predominantly in the right hemisphere. Scale bar: 100 µm.

**Supplementary Figure 7:**
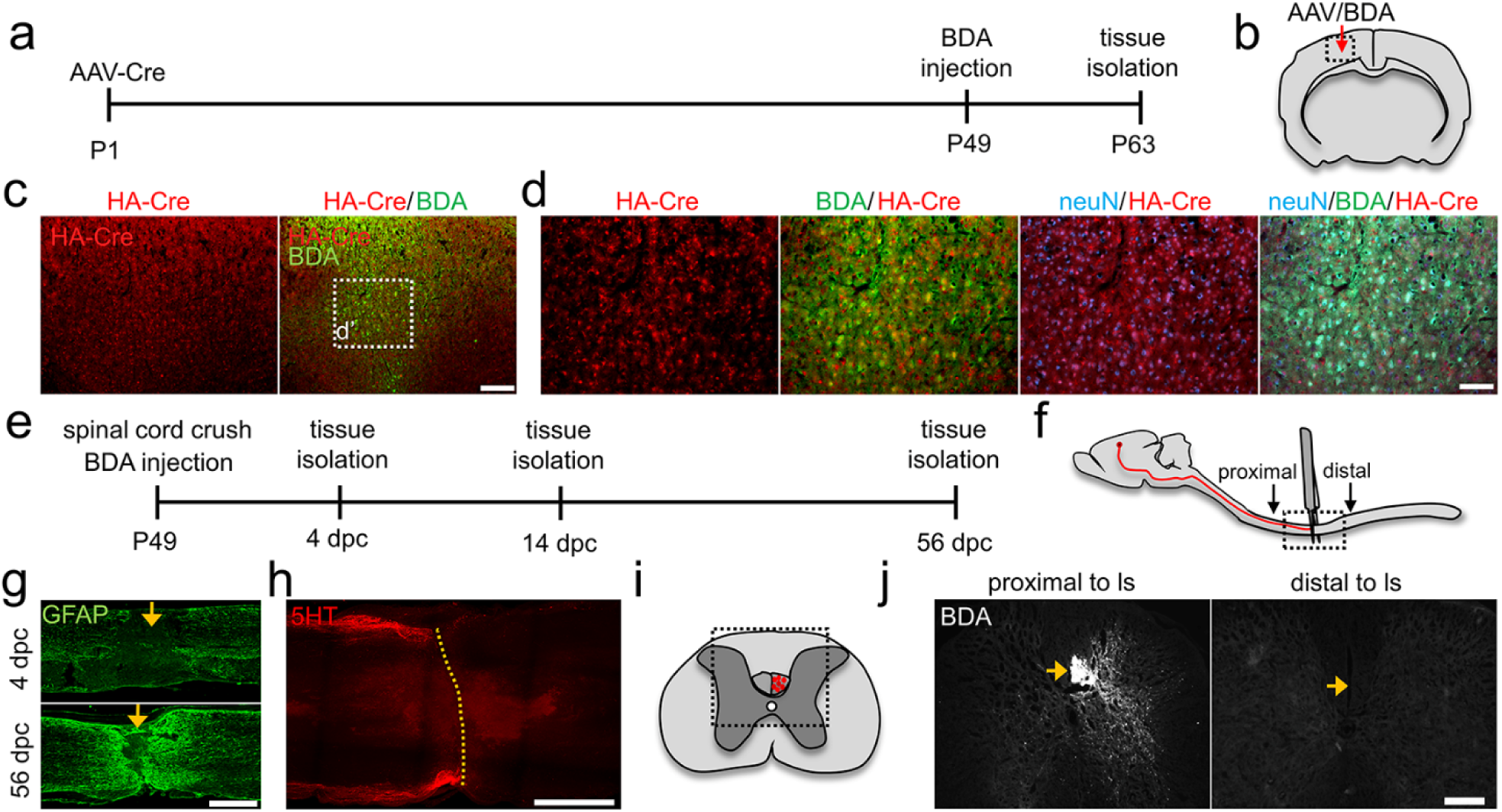
Validation of intracortical AAV2-application and spinal cord crush. **A)** Timeline for intracortical AAV2-HA-Cre (AAV2-Cre) treatment starting at postnatal day 1 (P1), BDA tracing of corticospinal neurons (P49), and tissue isolation (P63). **B)** Schematic drawing illustrates the virus and BDA injection site (arrow) in the sensorimotor cortex. The dashed box indicates the location of images presented in **C.** **C)** Coronal section of the sensorimotor cortex stained for the HA-Cre (red) and BDA (green). Dashed box indicates the magnified area in **D**. Scale bar: 200 µm. **D)** Higher magnification of the image shown in **C,** including NeuN staining (blue). Ca. 80% of BDA-positive neurons in layer V of the sensorimotor cortex were also HA-Cre-positive. Scale bar: 50 µm. **E)** Timeline for experiments verifying the completeness of spinal cord crush (SCC). P49 mice were subjected to SCC and tissues isolated 4, 14, or 56 days after spinal cord crush (dpc). **F)** Schematic drawing illustrating BDA tracing in the CST at the level of thoracic vertebra 8 (T8) and the SCC. The dashed box indicates the location of the immunohistochemical sections presented in **G and J**. **G)** Sagittal sections of thoracic spinal cord stained for GFAP (green) 4 and 56 dpc. Lack of astrocytes at the lesion site at 4 dpc verifies the completeness of the lesion. Eight weeks after crush, astrocytes bridged the lesion site. Scale bar: 500 µm. **H)** Maximum intensity projection of a confocal scan with a z-range of 500 µm from dorsal to ventral through a cleared spinal cord 4 dpc after prior serotonin (5-HT, red) immunostaining. Scale bar: 500 µm. **I)** Schematic drawing illustrating the location (dashed box) of images in **J** in spinal cord cross-sections. Labeled CST axons are indicated in red. **J)** Coronal sections of the spinal cord 2 weeks after SCC and intracortical BDA injection showing traced CST axons (arrow) proximal, but not distal to the lesion site (ls). Scale bar: 250 µm.

**Video S1: Open field locomotion after SCC and AAV2-GFP treatment**

The video shows a PTEN-floxed Ola mouse (PTEN^+/+^) that received an injection of AAV2-GFP into the left sensorimotor cortex at postnatal day 1. After 7 weeks, the mouse was subjected to complete spinal cord crush (SCC) (T8) and subsequently received an intracortical (left) injection of AAV2-GFP (see Fig. 1A). Videos were recorded at 1 and 8 weeks post SCC.

**Video S2: Open field locomotion after SCC and AAV2-hIL-6 treatment**

The video shows a PTEN-floxed Ola mouse (PTEN^+/+^) that received an injection of AAV2-GFP into the left sensorimotor cortex at postnatal day 1. After 7 weeks, the mouse was subjected to complete spinal cord crush (SCC) (T8) and subsequently received an intracortical (left) injection of AAV2-hIL-6 (see Fig. 1 A). Videos were recorded at 1 and 8 weeks post SCC.

**Video S3: Open field locomotion after PTEN^-/-^ and SCC**

The video shows a PTEN-floxed Ola mouse that received an injection of AAV2-Cre (PTEN^-/-^) into the left sensorimotor cortex at postnatal day 1. After 7 weeks, the mouse was subjected to complete spinal cord crush (SCC) (T8) and subsequently received an intracortical (left) injection of AAV2-GFP (see Fig. 1A). Videos were recorded at 1 and 8 weeks post SCC.

**Video S4: Open field locomotion after PTEN^-/-^ and SCC with AAV2-hIL-6 treatment**

The video shows a PTEN-floxed Ola mouse that received an injection of AAV2-Cre (PTEN^-/-^) into the left sensorimotor cortex at postnatal day 1. After 7 weeks, the mouse was subjected to complete spinal cord crush (SCC) (T8) and subsequently received an intracortical (left) injection of AAV2-hIL-6 (see Fig. 1A). Videos were recorded at 1 and 8 weeks post SCC.

**Video S5: Open field locomotion after DHT-mediated depletion of raphe spinal input**

The video shows a PTEN-floxed Ola mouse (PTEN^+/+^) that was subjected to complete spinal cord crush (SCC) (T8) and subsequently received bilateral intracortical injections of AAV2-hIL-6. To destroy raphe spinal input, the animal was injected intracerebroventricularly with the serotonin neurotoxin 5,7-dihydroxytryptamine (DHT) 6 weeks after SCC. Videos were recorded at 1 and 6 weeks post SCC and at 1d and 1 week after DHT application. Additionally, another mouse of the same background that received a similar treatment but was injected with AAV2-GFP instead of AAV-hIL-6 was recorded 6 weeks after SCC and 1 day after DHT treatment.

**Video S6: Open field locomotion determined in a BL6 mouse after SCC and AAV2-hIL-6 treatment**

The video shows a C57BL/6J wildtype mouse (BL6) that was subjected to complete spinal cord crush (SCC) (T8) and subsequently received an intracortical injection of AAV2-hIL-6 into the left sensorimotor cortex. Videos were recorded at 1 and 8 weeks post SCC.

**Video S7: Intracortical hIL-6 application induces STAT3 phosphorylation in raphe nuclei**

Three-dimensional projection of transverse confocal scan through 150 µm of cleared brain stem tissue (presented as projection in Fig. 7D) after unilateral (left) intracortical injection of AAV2-hIL-6 and BDA treatment. Serotonin (5-HT) was stained in blue, phosphorylated STAT3 (pSTAT3) in red, and BDA labeled CST axons in green.

## Acknowledgments

We would like to thank Marcel Kohlhaas and Anastasia Andreadaki for technical support and to Dr. Daniel Terheyden-Keighley for helpful comments on the manuscript. This work was supported by the German Research Foundation.

